# Genetic, vocal, and body size divergence across the Northern Peruvian Low supports two species within the Masked Flowerpiercer (*Diglossa cyanea*)

**DOI:** 10.1101/2022.05.18.492535

**Authors:** Silvia C. Martínez-Gómez, Carlos Esteban Lara, J. V. Remsen, Robb T. Brumfield, Andrés M. Cuervo

**Affiliations:** Instituto de Ciencias Naturales, Universidad Nacional de Colombia, Bogotá, Colombia; Dirección Académica Sede de La Paz, Universidad Nacional de Colombia, La Paz, Colombia; Louisiana State University, Museum of Natural Science, Baton Rouge, LA 70803, U.S.A

**Keywords:** Andes, biogeography, biological collections, *Diglossopis*, speciation, species delimitation, vocalizations, Andes, biogeografía, delimitación de especies, *Diglossopis*, especiación, vocalizaciones

## Abstract

Populations that become genetically isolated by geographical barriers may express phenotypic divergence more strongly in some traits than in others. Even when genetic differentiation among isolated populations accumulates at a rapid rate, this may not be reflected in phenotypic differentiation. This decoupling of trait divergence from genetic divergence has been found in multiple Andean bird lineages that occupy latitudinally long, linear ranges that are fragmented by ecological and topographic barriers. One of these montane birds is *Diglossa cyanea* (Thraupidae; Masked Flowerpiercer), a species with a distribution bisected by valleys and lowlands. Across these dispersal barriers one finds subspecies that differ only in subtle but diagnostic phenotypic differences. We evaluated genetic and phenotypic divergence throughout its distribution and found support for two distinct lineages sharply separated by the Marañón River valley at the Northern Peruvian Low (NPL). Specifically, we found that the two populations from the opposite sides of the NPL show deep divergence in mitochondrial DNA (mtDNA; ∼6.7% uncorrected *p* distance, n = 122), in song structure (exclusive final notes in southern populations, n = 88), and in wing length (longer wings in the southern population, n = 364). No genetic variation or song structure was observed within the large range of the southern group (from the NPL to central Bolivia) or within all northern populations (from the NPL to Venezuela). Moreover, these two lineages are possibly paraphyletic with respect to *D. caerulescens* (Bluish Flowerpiercer). Our results suggest that the southern taxon, *D. c. melanopis,* should be recognized as a species-level taxon, distinct from a redefined *D. cyanea*. We highlight the need to continue amassing complementary suites of datasets from field observations and experiments, laboratory analyses, and collection-based assessments, to better characterize the evolutionary history and taxonomic diversity of Neotropical montane birds.

**GRAPHICAL ABSTRACT:** 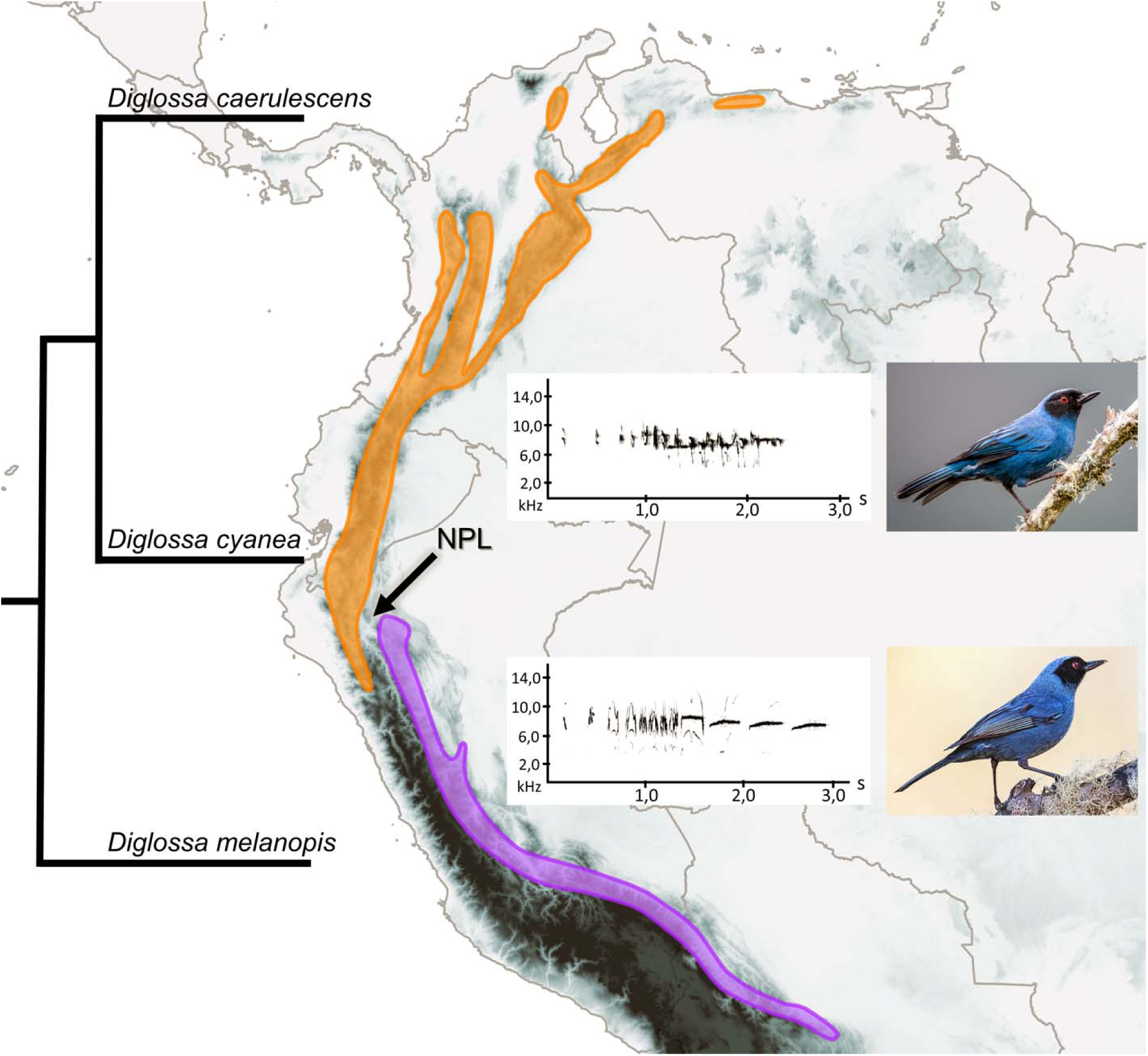

**LAY SUMMARY:**

⍰ We analyzed variation in genetics, songs, and body size among populations of a bird species of the Andean mountains (Masked Flowerpiercer) that shows relatively little plumage variation across its 3800-km long distribution from northern Venezuela to central Bolivia.
⍰ We found that the Masked Flowerpiercer consists of two divergent populations separated by a deep geographic depression known as the Northern Peruvian Low (NPL)
⍰ The degree of genetic divergence between them (in terms of mtDNA) is as great as between either of them and another species, the Bluish Flowerpiercer.
⍰ The two populations also have distinct songs, with the song of the southern group ending with clear whistles that are completely absent in the northern group. The southern group also tend to have longer wings than do populations north of the NPL.
⍰ The southern population should be treated as a separate species, *Diglossa melanopis*.

## RESUMEN

Las poblaciones que se encuentran genéticamente aisladas por barreras geográficas podrían expresar mayor divergencia fenotípica en algunos rasgos pero no en otros. Aún cuando la diferenciación genética entre poblaciones aisladas se obtenga rápidamente, esto podría no reflejarse en diferenciación fenotípica. Este desacople de divergencia ha sido observado en varios complejos de aves andinas que ocupan distribuciones latitudinalmente largas y fragmentadas por barreras ecológicas y topográficas. Una de estas aves es *Diglossa cyanea* (Thraupidae), un ave andina cuya distribución es fragmentada por enclaves secos, los cuales suelen estar asociados con límites de distribución de subespecies que difieren en rasgos fenotípicos apenas perceptibles. Evaluamos la divergencia tanto genética como fenotípica de *D. cyanea* a lo largo de su distribución y encontramos evidencia de dos linajes claramente separados por el cañón del río Marañón en la Depresión del Norte del Perú (DNP). Encontramos que las poblaciones a lados opuestos de la DNP muestran fuerte divergencia en ADN mitocondrial (∼6,7% distancia *p* no corregida, n = 122), en estructura del canto (con notas finales exclusivas de la población sur, n = 88) y en longitud de ala (alas más largas en población sur, n = 364). No observamos divergencia genética o de estructura del canto dentro de la amplia distribución del grupo sur (desde el DNP hasta Bolivia), ni entre poblaciones del grupo norte (desde el DNP hasta Venezuela). Adicionalmente, estos dos linajes son posiblemente parafiléticos con respecto a *D. caerulescens*. Nuestros resultados sugieren que el taxón sur, *D. c. melanopis* debe reconocerse como una especie distinta de *D. cyanea*. Destacamos la necesidad de continuar recolectando datos complementarios, incluyendo observaciones de historia natural y experimentos, análisis genómicos y colecciones biológicas para conseguir una mejor caracterización de la historia evolutiva y taxonómica de las aves montanas tropicales.

## INTRODUCTION

The degree of phenotypic divergence does not always parallel that of genetic divergence between separate populations. This decoupling between phenotypic trait differentiation and population genetic divergence may impede the characterization of biodiversity. Allopatric divergence driven by landscape changes or dispersal events is the most pervasive speciation mechanism underlying avian diversity (Graves 1985; Coyne and Orr 2004; Price 2008; Fjeldså et al. 2012). However, the extent of morphological, behavioral, and genetic divergence among daughter species may not be equally evident in all cases. Given sufficient time, the differentiation of geographically isolated populations accrues (Mayr 1942; Smith et al. 2014) at a pace that can be influenced by population size and natural selection (Nosil 2008; Winger and Bates 2015).

The speciation process in birds depends strongly on the interactions between topographic complexity, ecology, and dispersal ability, and the time required for divergence to act upon morphological, behavioral, and genetic traits (Price 2008; Benham et al. 2014; Smith et al. 2014). More specifically, the humid montane habitats of Andean birds have generated distributions that are fragmented by barriers of various sizes that could prevent gene flow, setting the stage for population divergence and, ultimately, speciation (Remsen 1984; Graves 1988, 1991; Weir 2009; Fjeldså et al. 2012). Studies on several widespread avian species complexes of the Andes that incorporate vocal data have demonstrated that species-level diversity is highly underestimated (Krabbe and Schulenberg 1997; Cadena and Cuervo 2010; Isler et al. 2020). This is because the tempo and magnitude of divergence in characters associated with reduction of gene flow are often decoupled in the speciation process (Mayr 1963; Winger and Bates 2015). For instance, marked phenotypic differences exist between recently diverged, almost genetically indistinguishable populations of *Coeligena* hummingbirds (Palacios et al. 2019). On the other hand, phenotypically indistinguishable populations with deep genetic divergences are found in several Andean bird complexes, including ducks (Gutiérrez-Pinto et al. 2019), hummingbirds (Chaves et al. 2011; Benham et al. 2014), suboscines (Valderrama et al. 2014; Cadena et al. 2020), and oscines (Gutiérrez-Pinto et al. 2012; Prieto-Torres et al. 2018; Cadena et al. 2019). That is, true diversity may be frequently overlooked because characterizations based on genetic or phenotypic traits alone are often insufficient.

Tanagers in the flowerpiercer clade *Diglossa* (Thraupidae) are ecologically and phenotypically specialized members of Neotropical montane bird communities (Moynihan 1968; Vuilleumier 1969), with a peak of diversity in the Andes. With 18 recognized species (Bock 1985; Isler and Isler 1999; Dickinson and Christidis 2014), *Diglossa* is one of the genera encompassing Andean birds that has revealed some of the most extraordinary and intriguing leapfrog patterns of geographic variation and taxonomic bias in birds (Moynihan 1979; Graves 1982; Vuilleumier 1984; Mauck and Burns 2009). For example, molecular data have shown a rapid diversification in the core *Diglossa* group (Mauck and Burns 2009), with a much more convoluted history than previously thought (see Gutiérrez-Zuluaga et al. 2021). The most morphologically similar taxa may not be sister lineages, implying a complex genetic basis of divergence yet to be revealed (Hiller et al. 2021). In contrast to the core *Diglossa* group, the three species in the *Diglossopis* clade (*D. caerulescens*, *D. cyanea,* and *D. glauca*; Mauck and Burns 2009) show deeper divergences but only modest phenotypic differences. This group consists of similar-looking flowerpiercers with bluish plumages, facial masks, and reduced bill hooks; they are largely sympatric along the tropical Andes, and are found across major topographic discontinuities such as the Marañón valley.

The Northern Peruvian Low (NPL) is the preeminent Andean barrier, largely defined by low summits and deep dry canyons (Vuilleumier 1968, 1984), especially the Porculla Pass, the Huancabamba Depression, and the Marañón river valley in northern Peru (Duellman 1979; Parker et al. 1985; Cuervo 2013). The NPL is well known to be a barrier that shapes geographic range limits (Vuilleumier 1968; Cracraft 1985) and gene flow of Andean birds (Cuervo 2013), especially for those adapted to humid and forested ecosystems, as in most *Diglossa*. However, not all taxa are equally affected by this geographic break (Vuilleumier 1984; Parker et al. 1985; Weir 2009; Cuervo 2013; Winger and Bates 2015), including *Diglossa*.

The Masked Flowerpiercer is abundant and widespread in the tropical Andes where it occurs in montane and stunted forests, semi-open areas with isolated trees and scrubs, and forest borders. Although spanning a wide latitudinal range from northern Venezuela to central Bolivia (∼4,500 km), its elevational range is restricted to 1,800-3,600 m. a.s.l. (Fjeldså and Krabbe 1990; Parker et al. 1996), which often amounts to only a few kilometers in linear distance. Five subspecies are currently recognized (Dickinson and Christidis 2014), from south to north: *melanopis, dispar, cyanea, obscura*, and *tovarensis*. These taxa were defined on the basis of subtle differences in size or plumage color (Hellmayr 1935; Zimmer 1942; Meyer de Schauensee 1951; Zimmer and Phelps 1952; Isler and Isler 1999). Range boundaries between the five subspecies are clearly delimited by geographic features, except for *D. c. cyanea* and *D. c. dispar*, which replace each other in southern Ecuador (Fjeldså and Krabbe 1990; Freile and Restall 2018) (Figure 1A). Although only vague characterization exists of the intraspecific phenotypic differences for northern *D. cyanea* populations, implying they may not be phenotypically diagnosable units, the small differences between northern *D. cyanea* subspecies that have been described include subtleties in coloration and facial masks (Zimmer and Phelps 1952), and vocal peculiarities in *D. c. tovarensis*, albeit with small sample sizes (Fjeldså and Krabbe 1990; Hilty 2003). The subspecies *D. c. dispar* is described to be similar in size to *D. c. cyanea*, but similar in plumage to *D. c. tovarensis* (Zimmer and Phelps 1952). These differences between *D. cyanea* subspecies are slight and were described on the basis of a handful of specimens for comparisons (Vuilleumier 1969); hence, whether these recognized taxa are diagnosable units need to be addressed (Patten 2010; Remsen 2010). In contrast, the subspecies *D. c. melanopis* from south of the NPL in Peru south to Bolivia (Schulenberg et al. 2010; Herzog et al. 2017) is more clearly defined. This southern subspecies is larger and duller, with a paler forecrown and less prominent white tips on the undertail coverts compared to the other four subspecies from north of the NPL (Hellmayr 1935; Isler and Isler 1999).

**Figure 1.**
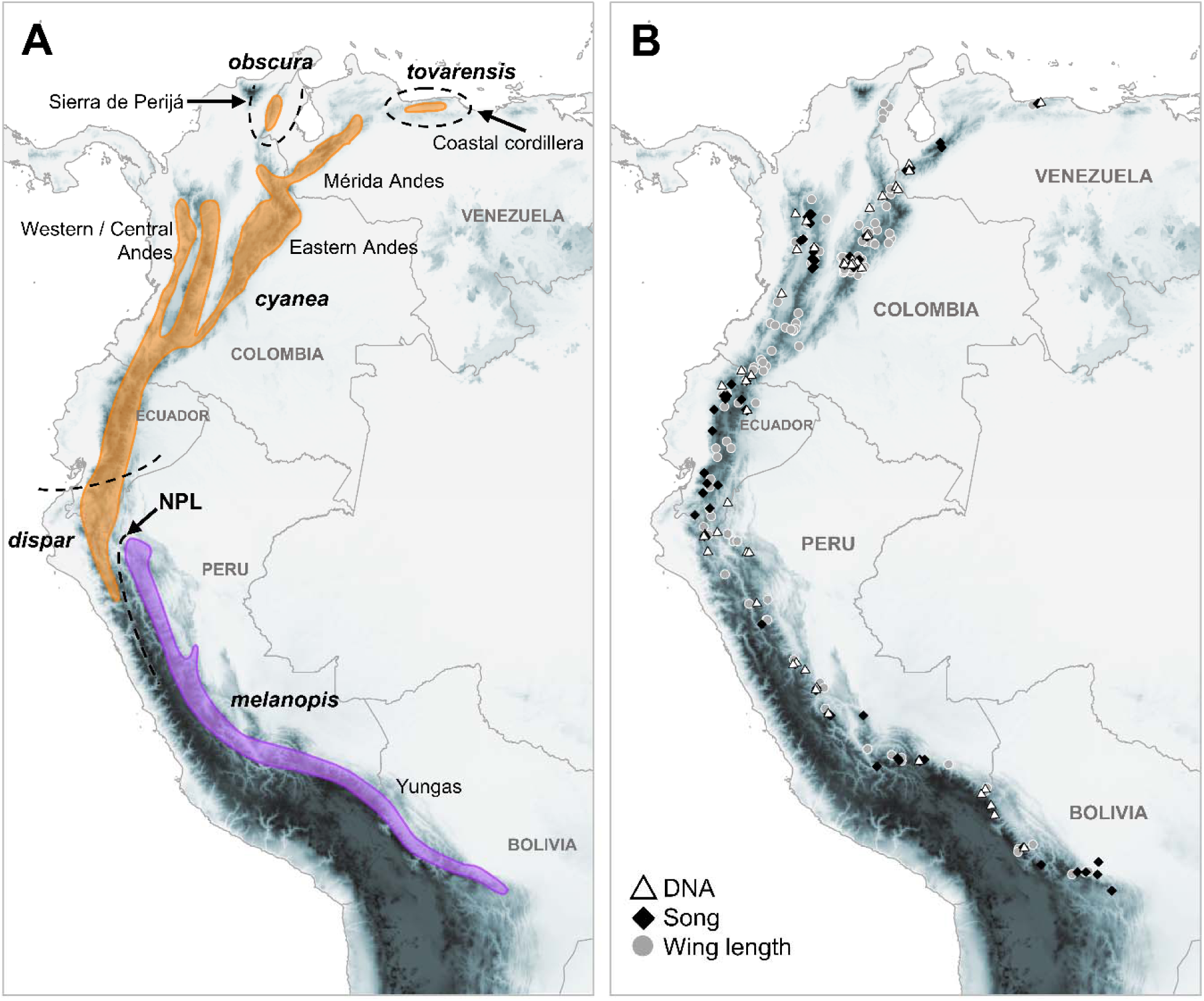
(A) Geographic distribution of the Masked Flowerpiercer, *Diglossa cyanea*. Dashed black lines indicate approximate range boundaries between the five subspecies currently recognized. Note that except for the northern limit of *D. c. dispar*, subspecies boundaries coincide with low elevation gaps. The two colors denote the northern (orange) and southern (purple) groups separated by the Northern Peruvian Low (NPL). (B) Symbols represent sample localities for vouchered specimens of *D. cyanea* used in the phylogeographic analysis (white triangles), wing length from study skins (gray dots), and song data (black diamonds). We were unable to obtain vocal and genetic samples for *D. c. obscura*.

Here, we conducted a geographically comprehensive analysis of genetic variation integrated with vocal and morphological data to assess (1) how the Northern Peruvian Low (NPL) bisects populations of *D. cyanea*; and (2) whether genetic structure, vocal structure, and body size correspond to the current subspecies taxonomy. First, we explored the phylogeographic patterns of *D. cyanea* to evaluate the extent of its genetic structure in a geographic context. We discovered that one of the two major clades of *D. cyanea* seems closer to *D. caerulescens*; thus, we expanded the geographic sampling of molecular data in that species. Second, we compared song structure between the two *D. cyanea* groups revealed by the genetic analysis. Third, we quantified differences in wing length, as an index of body size, between the *D. cyanea* groups.

## METHODS

### Phylogeographic Structure

Mauck and Burns (2009) found that *Diglossa cyanea* (Masked Flowerpiercer) and *D. caerulescens* (Bluish Flowerpiercer) are sister species; this would be one of the few cases in Andean birds of sister pairs that overlap for most of their geographic distributions. However, their analyses included only a single sample per species. Therefore, we expanded the geographic sampling to encompass the entire range of *D. cyanea* and, to a lesser extent, of *D. caerulescens.* To do this, we sequenced the mitochondrial gene NADH dehydrogenase subunit 2 (ND2, 1041 bp) for 122 individuals of *D. cyanea* and 33 of *D. caerulescens*. We included the samples used by Mauck and Burns (2009) for both species. We included three outgroups: *D. glauca*, the other species in the *Diglossopis* clade, and *D. albilatera* and *D. indigotica,* two species of the core *Diglossa* clade (see Supplementary Material Table S1 for details).

DNA extraction, amplification, and sequencing protocols followed Cuervo et al. (2014). Raw sequence data were inspected for ambiguities and stop codons using Sequencher 4.7 (GeneCodes Corp., Ann Arbor, MI), and were aligned using Geneious 9.1.8. We estimated ND2 gene trees using RAxML 8.2.12 (Stamatakis 2014) and MrBayes 3.2.7a (Ronquist et al. 2012) via the CIPRES Science Gateway 3.3 portal (Miller et al. 2010). For RAxML, we implemented the GTRCAT approximation for rate heterogeneity with 25 distinct categories, and automatic rapid bootstrapping search to assess nodal support after 650 replicates with the autoMRE option. For MrBayes, we implemented a partition scheme with a model of substitution for each codon position (first: HKY + Γ, second: HKY + I, third: GTR + I + Γ) as suggested by the Akaike Information Criterion with correction (AICc) in PartitionFinder 2 (Lanfear et al. 2017). We ran four Markov Chain Monte Carlo (MCMC) chains for 20 million generations, sampling every 1,000^th^, discarding the initial 50% as burn-in. We also used BEAST 2.6.3 (Bouckaert et al. 2019) to estimate divergence times while simultaneously estimating a tree topology (Heled and Drummond 2010). We applied a lognormal relaxed clock and used the default Yule Process as the tree prior. We used the average ND2 substitution rate (2.5% per million years) estimated for other tropical passerine birds (Smith and Klicka 2010). The alignment contained 158 taxa, 1041 sites, and 256 variable sites. We ran two independent MCMC runs starting from random trees for 100 million generations, sampling every 5,000^th^ and discarding the first 50% as burn-in in each run. Both posterior parameter values and tree files were combined with resampling in LogCombiner to obtain 10,002 post-burning parameter estimates and trees, respectively. We used TreeAnnotator 2.6.3 (Bouckaert et al. 2019) to calculate a maximum clade credibility (MCC) tree with mean heights. We inspected convergence in the post burn-in MCMC parameter estimates for both Bayesian analyses in Tracer 1.7.1 (Rambaut et al. 2018). To further examine the genetic divergence between subspecies and major groups, we estimated genetic distances in MEGA (Kumar et al. 2018; Stecher et al. 2020) and built median-joining networks with ε = 0 (Bandelt et al. 1999) for each major clade in PopART 1.7 (Leigh and Bryant 2015).

### Vocal Variation

We examined audio recordings of *D. cyanea* archived in the Macaulay Library (ML; Cornell Lab of Ornithology, Ithaca, NY) and xeno-canto (XC; www.xeno-canto.org) (Supplementary Material Table S2, Figure 1). We retrieved a total of 88 recordings of *D. cyanea,* distributed by phylogeographic groups as 55 north of the NPL and 35 south of the NPL, and distributed by subspecies as follows: 35 of *D. c. melanopis*, 4 of *D. c. dispar*, 47 of *D. c. cyanea*, and 2 of *D. c. tovarensis*. Recordings of *D. c. obscura* were unavailable. We excluded short or fragmented songs, or recordings with low signal-to-noise levels. Our sampling unit for the vocal analysis (see below) was at the individual level. We assumed that each audio recording from the same locality and date belonged to the same individual. We analyzed up to 10 song bouts (mean _[song bouts]_ = 4.32, SD = 2.81) of each recording to capture the intra-individual variation. We defined song bouts in recordings as clusters of vocal elements exceeding a duration of 0.5 s, and separation from other clusters was distinguished by silence intervals exceeding 1 s.

Songs of *D. cyanea* consist of a high-pitched, melodic warble of complex, accelerating tweet notes. To fully explore vocal variation, we analyzed the full song and up to three divided sections, as follows (also see Figure 2): (1) the full song is equivalent to the total length of the song (hereafter F); (2) we separately analyzed the first section of the song (hereafter S1), which corresponds to the introductory phrase, consisting of 2–5 short (< 0.1 s) “*tzi*” notes, delivered at 0.1–0.4 s intervals; (3) the second section (hereafter S2) corresponds to the main phrase of the song: a complex, fast chatter of rich elements delivered at < 0.08 s intervals; and (4) a third section of the song (hereafter S3), when present, is a series of 3–4 clear but strident whistling notes.

**Figure 2.**
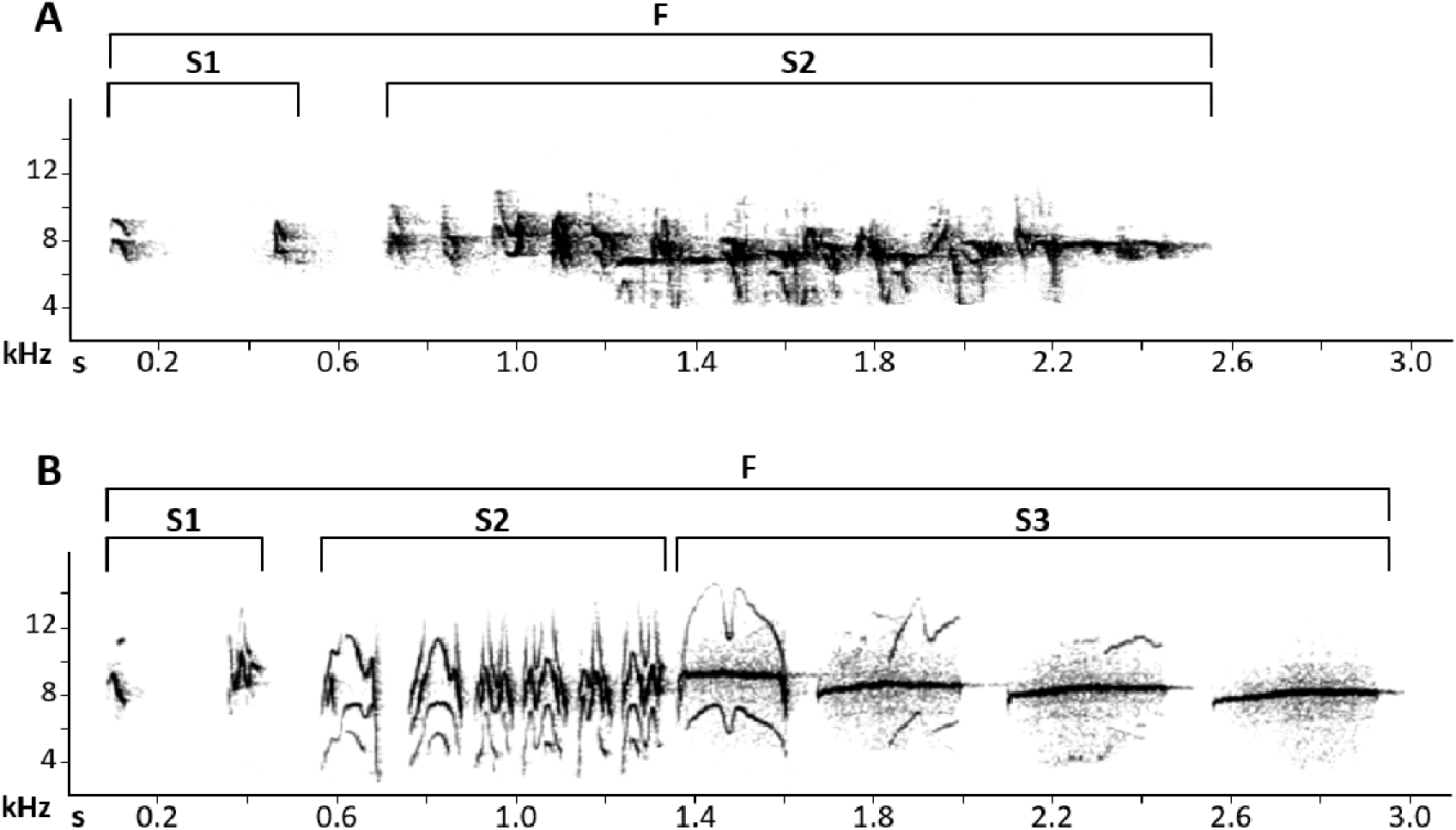
Spectrograms of representative samples of the typical song in the Masked Flowerpiercer, *D. cyanea.* (A) an example from the northern group (*D. cyanea cyanea* from Caldas, Colombia, XC-373183), (B) an example from the southern group (*D. c. melanopis* from Cochabamba, Bolivia, ML-87666). Phrase sections (S1, S2, S3), and the full song (F) are indicated by brackets. The striking difference in the typical song between the northern (A) and southern (B) groups of *D. cyanea* is driven by the shorter warbling chatter (S2) and presence of the terminal phrase of strident whistle notes (S3) in the southern group songs.

We measured five spectral and temporal traits in the full song (F) and each composing section (S1, S2, S3) by placing landmark boxes on spectrograms using Raven Pro 1.6 (Cornell Lab of Ornithology, Ithaca, NY), with custom visual settings (Hann type spectrograms; window size = 512 samples). These traits were: duration, peak frequency, maximum frequency, minimum frequency, and bandwidth. Upon examination of all available vocal samples, we identified two clear song types segregated geographically, and a much less frequent alternative song. The latter consists of a single, main unit similar in structure to S2 of the typical song (Supplementary Material Figure S1).

To assess quantitative differences in songs between the southern and northern groups, and between the four sampled subspecies (*D. c. melanopis*, *D. c. dispar*, *D. c. cyanea*, and *D. c. tovarensis*), we fitted multiple Bayesian Linear Mixed Models (BLMMs) using Markov chain Monte Carlo techniques with the *MCMCglmm* package (Hadfield 2010) implemented in R 4.0.1 (R Core Team 2021). We used the Gaussian error distribution and used *MCMCglmm* default settings and priors. We report standardized estimates of regression coefficients as the mean 1,000 posterior distributions with 95% credible intervals (CIs) and considered effects to be statistically significant if CIs did not overlap with zero. Specifically, to examine vocal differences between the southern and northern groups, we fit 15 BLMMs as follows (also see supplementary Material Table S3, Figure 1): we first used the full song (F) and fitted, separately, five models, one per each vocal trait (duration, peak frequency, maximum frequency, minimum frequency, and bandwidth), which were used as response variables, and the geographical group as a binary fixed effect. We then fit five models for the phrase section S1 and five models for the section S2, using the same structure, to total 15 models. To incorporate intra-sample variation (Bolker et al. 2009), we included the recording ID as a random effect in all models. Importantly, the response variables were centered and standardized, using the *scale* function in R, to have a mean = 0, and standard deviation = 1, which allowed us to compare effect size across models (Nakagawa and Cuthill 2007; Schielzeth 2010). Following the same settings and model structure as described above, we fit another 15 models (see Supplementary Material Table S3), but instead of using group as a binary fixed effect, we used subspecies as a categorical fixed effect. We used the *relevel* function in R to obtain the statistical differences between pairs of subspecies in each model.

### Body Size Variation

We examined body size variation of *D. cyanea* along its latitudinal distribution (Figure 1) using wing length (WL) as a surrogate for body size (Zink and Remsen 1986). Wing length may have problems in indexing body size for migratory birds where wing length varies in relation to the migratory activity, but for *Diglossa*, which are nonmigratory, that is not the case (Remsen, unpublished data). We measured WL (unflattened wing chord) using an end-stopped metallic ruler (± 1.0 mm) on 364 round skins: 125 *D. c. melanopis,* 19 *D. c. dispar,* 207 *D. c. cyanea,* four *D. c. tovarensis* and nine *D. c. obscura* (Supplementary Material Table S4). Also, we reviewed specimen records from museums, ornithological datasets, and the literature (e.g. Paynter 1981; Stephens and Traylor 1983; Paynter 1992; Núñez-Zapata et al. 2016) to georeference localities of historical specimens. We only considered specimens categorized as adults without evidence of molt or worn primaries.

To examine variation in WL along the latitudinal distribution of the species (17.24°S to 10.41°N), we fit linear models (LM) in R (R Core Team 2021). On a first LM, we only included data from the southern group and used latitude and its interaction with sex as predictors (to obtain the slope for each sex) and WL as the response variable (Wing Length ∼ Latitude*Sex). On a second LM, we used a similar model structure, but only included data from the northern group. Then, we fit two additional LMs to quantify statistical differences between the WL of males from the southern group vs. the northern group, and between the WL of females from the southern group vs. the northern group (Wing Length ∼ Geographical Group for males, and Wing Length ∼ Geographical Group for females).

## RESULTS

### The Northern Peruvian Low (NPL) divides *Diglossa cyanea* into two divergent lineages

The samples of *D. cyanea* clearly fell into one of two clades that are sharply associated with geography (Figure 3). The uncorrected pairwise genetic distance between these two groups averaged 6.7% (Table 1). The northern group contained all *D. cyanea* samples to the west and north of the NPL from Cajamarca, Peru, through the Northern Andes up to Aragua, in the Coastal Cordillera in northern Venezuela. The low genetic variation within this group (average pairwise base differences = 0.0028) implies little apparent structure associated with geography and a lack of differentiation in this marker among the northern subspecies (*D. c. dispar, D. c. cyanea, D. c. tovarensis*). All samples to the south and east of the NPL from Amazonas, Peru, south to Bolivia formed the second major clade in *D. cyanea*, corresponding entirely to the southern subspecies (*D. c. melanopis*). Genetic diversity within this southern group was larger than within the northern group, with two groups of haplotypes present in its southern range (Cusco and Puno in Peru, and La Paz in Bolivia), and a third haplotype group including all samples from central and northern Peru (Pasco, Huánuco, San Martín, and Amazonas).

**Table 1.** Pairwise genetic divergence between the *D. cyanea* named taxa and *D. caerulescens* (last six taxa; marked with asterisk). The southern *D. cyanea* group (S) is solely represented by *D. c. melanopis*, whereas the northern group (N) is represented by three out of four named taxa (*D. cyanea obscura* was not sampled). Upper right cells contain the average number of base differences per site between groups (uncorrected p-distance), with the pairwise deletion option. Lower left cells show the net average differences per site between groups. Diagonal cells contain the within-group average number of base differences per site.

| Subspecies | *Diglossa cyanea* |  |  |  | *Diglossa caerulescens* * |  |  |  |  |  |
| --- | --- | --- | --- | --- | --- | --- | --- | --- | --- | --- |
|  | *tovarensis* (N) | *cyanea* (N) | *dispar* (N) | *melanopis* (S) | *caerulescens* | *ginesi* | *saturata* | *ssp.* | *media* | *mentalis* |
| *tovarensis* (N) | - | 0.0071 | 0.0077 | 0.0706 | 0.0740 | 0.0754 | 0.0740 | 0.0780 | 0.0730 | 0.0786 |
| *cyanea* (N) | 0.0058 | 0.0025 | 0.0029 | 0.0667 | 0.0740 | 0.0754 | 0.0740 | 0.0780 | 0.0729 | 0.0783 |
| *dispar* (N) | 0.0067 | 0.0007 | 0.0020 | 0.0673 | 0.0744 | 0.0755 | 0.0744 | 0.0784 | 0.0734 | 0.0788 |
| *melanopis* (S) | 0.0670 | 0.0618 | 0.0626 | 0.0073 | 0.0799 | 0.0813 | 0.0804 | 0.0822 | 0.0788 | 0.0837 |
| *caerulescens* * | 0.0740 | 0.0727 | 0.0734 | 0.0762 | - | 0.0034 | 0.0043 | 0.0156 | 0.0106 | 0.0138 |
| *ginesi* * | 0.0749 | 0.0737 | 0.0740 | 0.0772 | 0.0029 | 0.0010 | 0.0044 | 0.0170 | 0.0120 | 0.0152 |
| *saturata* * | 0.0716 | 0.0703 | 0.0710 | 0.0744 | 0.0019 | 0.0015 | 0.0048 | 0.0167 | 0.0117 | 0.0149 |
| *ssp.* * | 0.0778 | 0.0766 | 0.0772 | 0.0783 | 0.0154 | 0.0163 | 0.0141 | 0.0004 | 0.0108 | 0.0159 |
| *media* * | 0.0730 | 0.0716 | 0.0724 | 0.0752 | 0.0106 | 0.0115 | 0.0093 | 0.0106 | - | 0.0070 |
| *mentalis* * | 0.0779 | 0.0763 | 0.0771 | 0.0794 | 0.0131 | 0.0140 | 0.0118 | 0.0150 | 0.0063 | 0.0015 |

**Figure 3.**
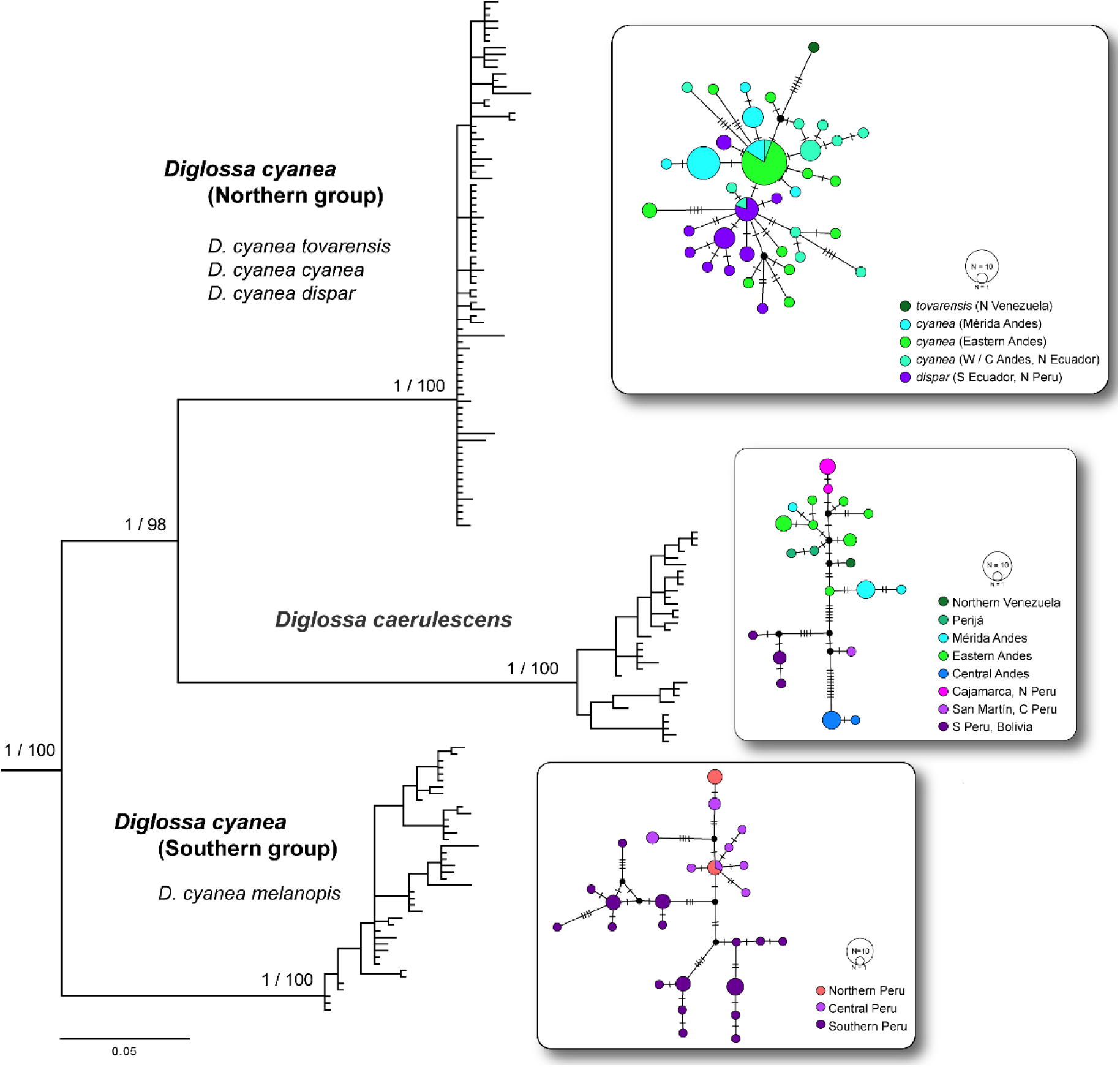
The ND2 gene tree is a 50% majority-rule consensus tree from MrBayes showing the three major clades corresponding to the northern (top, n = 81) and southern (bottom, n = 41) populations of Masked Flowerpiercer, *D. cyanea*, geographically divided by the Northern Peruvian Low, and multiple samples of the Bluish Flowerpiercer, *D. caerulescens* (middle, n = 33). Values on branches indicate nodal support as Bayesian posterior probability and maximum likelihood bootstrap support. In front of each major clade, median-joining haplotype networks depict genetic diversity and relationships among haplotypes within groups. Color denotes geographic regions or subspecies as currently defined.

Furthermore, the estimated ND2 tree showed that the northern and southern clades *of D. cyanea* were not sister to each other: all the *D. caerulescens* samples formed a clade that was sister to the northern group of *D. cyanea* in all analyses. Uncorrected pairwise genetic distances between *D. caerulescens* and the northern and southern groups of *D. cyanea* averaged 7.5% and 8.1%, respectively (Table 1). We did not aim to sample *D. caerulescens* in detail, but we found genetic structure associated with geography and the subspecies taxonomy that describes its phenotypic diversity (Supplementary Material Figure S2). We estimated that the two deepest divergence events leading to each of the three major clades (i.e., southern *D. c. melanopis*, *D. caerulescens*, and northern *D. cyanea*) occurred rapidly during the Pliocene (3.7–3.0 million years, Mya), and differentiation within ocurred in the Pleistocene (0.58–0.25 Mya, Supplementary Material Figure S3).

### Two Song Types in *Diglossa cyanea* Reflect the Genetic Structure

We found two general song types that separate *D. cyanea* into a northern and a southern group across the NPL, reflecting the phylogeographic results. Songs clearly differ in note structure, duration, and spectral metrics between both sides of the NPL (Figure 4). Specifically, songs of the southern group (*D. c. melanopis*) are characterized by the addition of a terminal series of strident whistling notes (S3) that are completely absent from our sample of recordings of the northern group (*D. c. tovarensis*, *D. c. cyanea*, and *D. c. dispar*); to the best of our knowledge, this terminal series has never been reported in field observations of *D. cyanea* north of the NPL. Additionally, the southern songs have a much shorter (β_southern_ = 0.7 s, 95% Cis = 0.5 s to 0.8 s) warbling chatter (S2) than songs of the northern group (β_northern_ = 1.9 s, 95% Cis = 1.8 s to 1.9 s, contrast β_southern_–β_northern_ = 1.2 s, 95% Cis = 1.0 s to 1.3 s; see details in Supplementary Material Table S5 and S6). Although lacking the terminal whistling phrase (S3), songs of the northern group have longer phrases (S1, S2), yielding no statistical difference in full song duration (F) between groups. Moreover, songs of the southern group (*D. c. melanopis*) are higher pitched (i.e., greater peak frequency values across comparable song sections; contrast β_northern_–β_southern_ = 1004 Hz, 95% Cis = 860 Hz to 1156 Hz), and tend to occupy narrower bandwidths due to their higher minimum frequencies than the northern songs (bandwidth contrast β_southern_–β_northern_ = 1352 Hz, 95% Cis = to 902 Hz to 1785 Hz; see greater detail in Supplementary Material Table S5 and S6).

**Figure 4.**
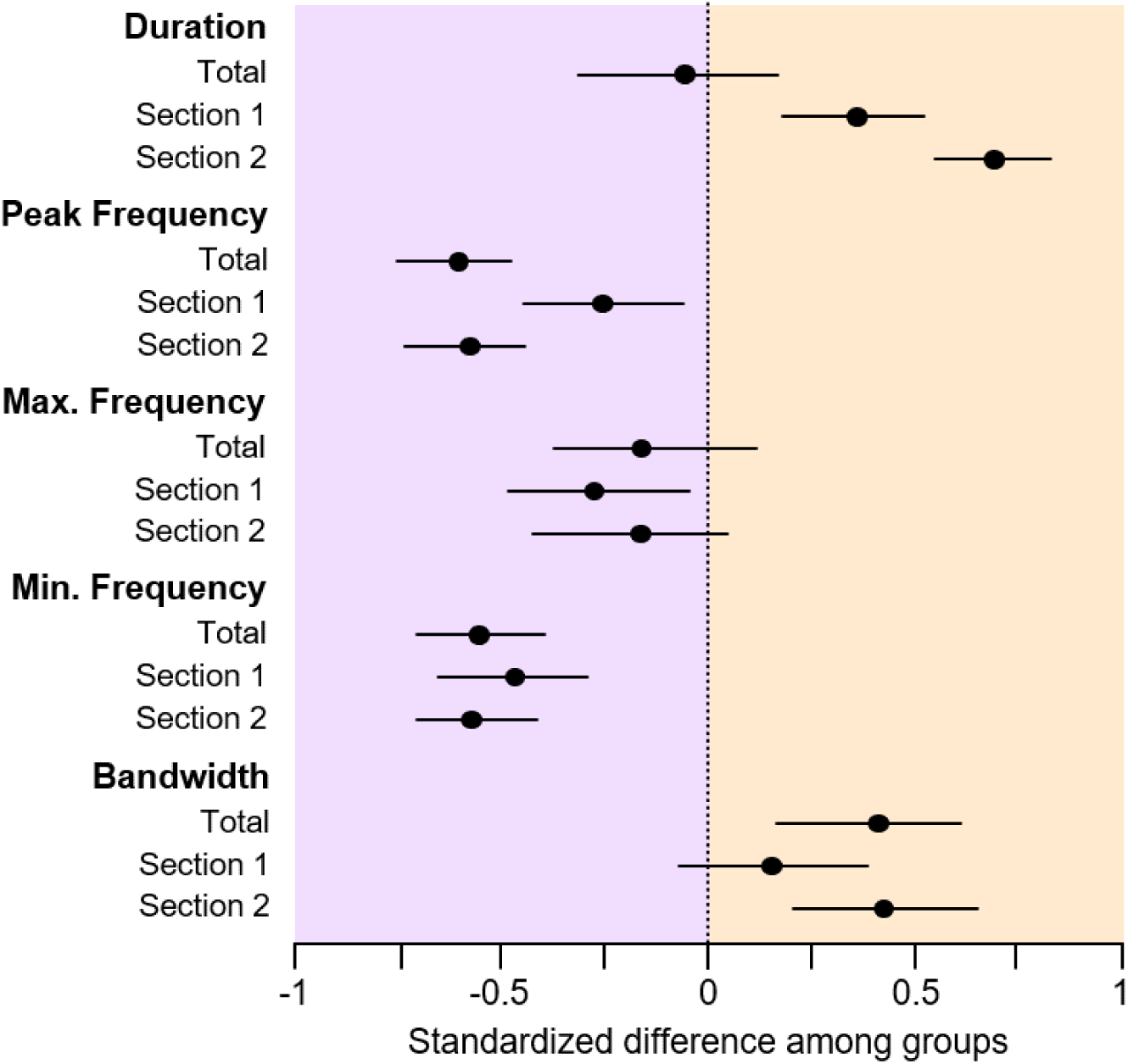
Acoustic differences and their confidence intervals (horizontal lines) in the songs of the two *D. cyanea* groups, separated by the Northern Peruvian Low (NPL). Differences were calculated as the mean values of the northern vocal samples minus those of the southern samples, where a standardized difference of 0 indicates no difference between the means of the two groups, positive standardized differences indicate larger values in the northern group (north and west of the NPL to Venezuela), and negative standardized differences indicate larger values in the southern group (*D. c. melanopis* from the east and south of the NPL). Although full song duration was similar between the two groups, songs of southern groups were higher pitched with narrower bandwidths.

To further explore vocal differences among the currently defined subspecies of *D. cyanea*, we compared the five vocal variables in the full song and the two shared sections (F, S1, S2) by considering each of the northern subspecies (*D. c. tovarensis*, *D. c. cyanea*, *D. c. dispar*) separately, except for *D. c. obscura* for which no vocal data exist. The southern subspecies (*D. c. melanopis*) differed vocally from the other *D. cyanea* subspecies of the northern group, particularly for the higher pitch of its song elements, revealed by higher values of peak frequency and minimum frequency; see Fig. 5, details in Supplementary Material Table S7). Lastly, the northern subspecies (*D. c. tovarensis*) has a shorter full song (contrast β*_tovarensis_*–β*_dispar_* = 0.9 s, 95% Cis = 0.2 s to 1.5 s), accompanied by a much lower maximum frequency in S1 and S2 overall, and outstanding visual characteristics of the notes in comparison to other northern *D. cyanea* subspecies, although sample sizes are limited (n = 2) (Supplementary Material Table S7, Supplementary Material Figure S4).

**Figure 5.**
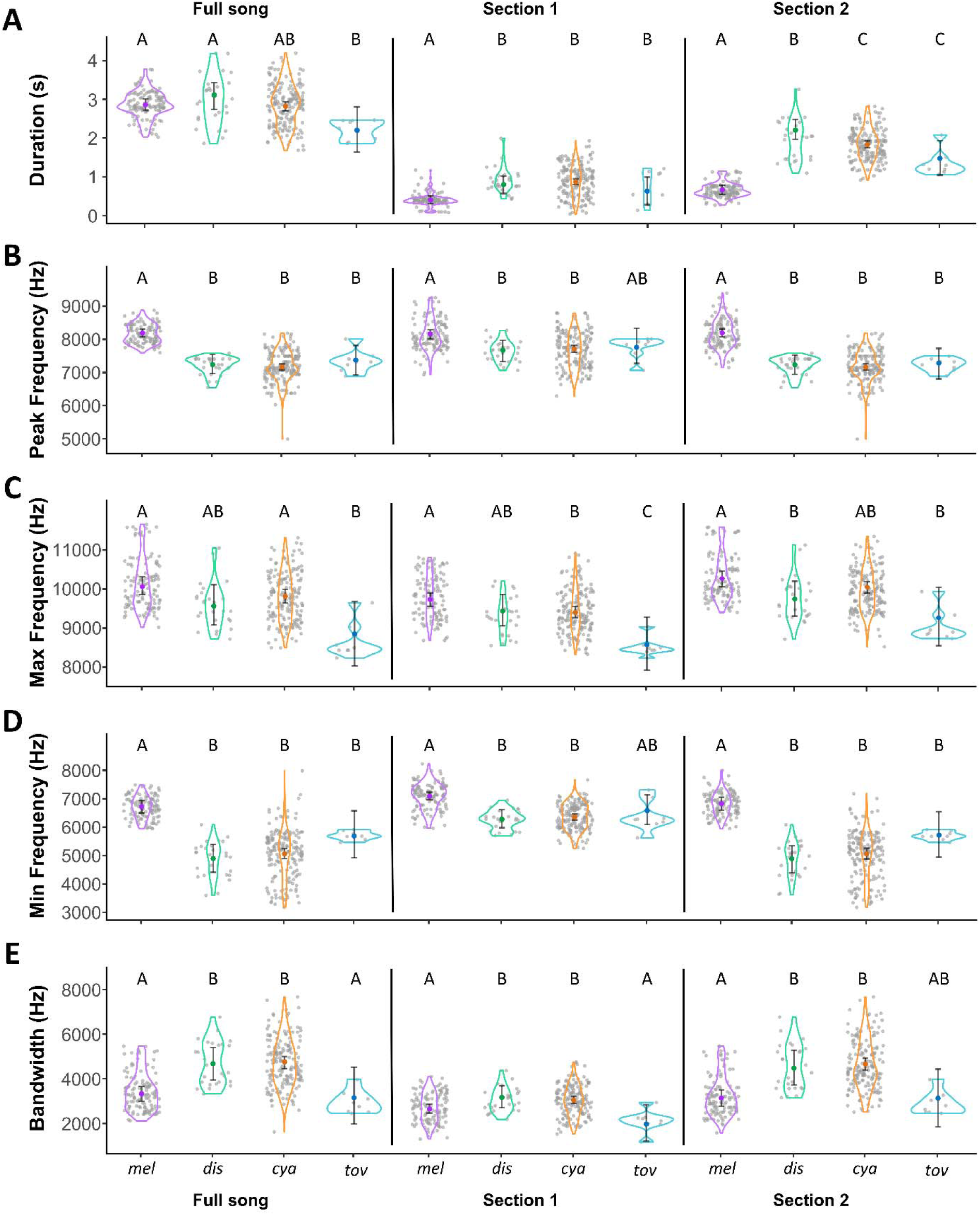
Quantitative acoustic differences among four subspecies of *D. cyanea* (all except the unsampled *obscura*); *mel*: *melanopis* (southern group); *dis*: *dispar*, *cya*: *cyanea*, *tov*: *tovarensis* (northern group). Acoustic variables analyzed included (A) duration, (B) peak frequency, (C) maximum frequency, (D) minimum frequency, and (E) frequency bandwidth. The full song and phrase sections 1 and 2 are compared among taxa. In most cases, the southern subspecies *D. c. melanopis* showed the most dissimilar acoustic traits, being most different to songs of the northern subspecies *D. c. cyanea*.

In addition to the song differences between the two *D. cyanea* groups from opposite sides of the NPL, we found that in about half of the available recordings of the southern group (*D. c. melanopis*), an alternative, or secondary song was present (see Supplemental Material Figure S1). The alternative song consists of a single, continuous chatter phrase, largely different in structure from the typical song on either side of the NPL, characterized by a slightly shorter duration (β_alternative_ = 2.5 s, 95% CI = 2.2 s to 2.7 s; β_southern_= 3.0 s, 95% CI = 2.8 s to 3.1 s) and lower frequencies (both peak and minimum) than the typical southern song (Supplemental Material Table S8).

### Wing length (WL) differs between the southern and northern groups

We only found a statistically significant (and positive) association between latitude and wing length (WL) for females in the southern group (β_females southern_ = 0.25, p = 0.01, Figure 6A, Supplementary Material Table S9). Wings of *D. c. melanopis* females tended to be longer towards its northern range limit. In sharp contrast, neither WL of males nor females of the northern group showed any statistically significant variation along latitude (i.e., confidence intervals of slopes included 0, β_males southern_ = 0.099, p = 0.34, β_females northern_ = 0.082, p = 0.37, β_males northern_ = −0.003, p = 0.98). More importantly, we found that WL differed statistically between the two groups across the NPL for both, females (β_females southern_ – β_females northern_ = 3.8 mm, p<0.001) and males (β_males southern_ – β_males northern_ = 4.8 mm, p<0.001, Figure 6B, Supplementary Material Table S10). Simply put, southern females have longer wings than do northern females, and southern males have longer wings than do northern males (Figure 6B, Supplementary Material Table S10).

**Figure 6.**
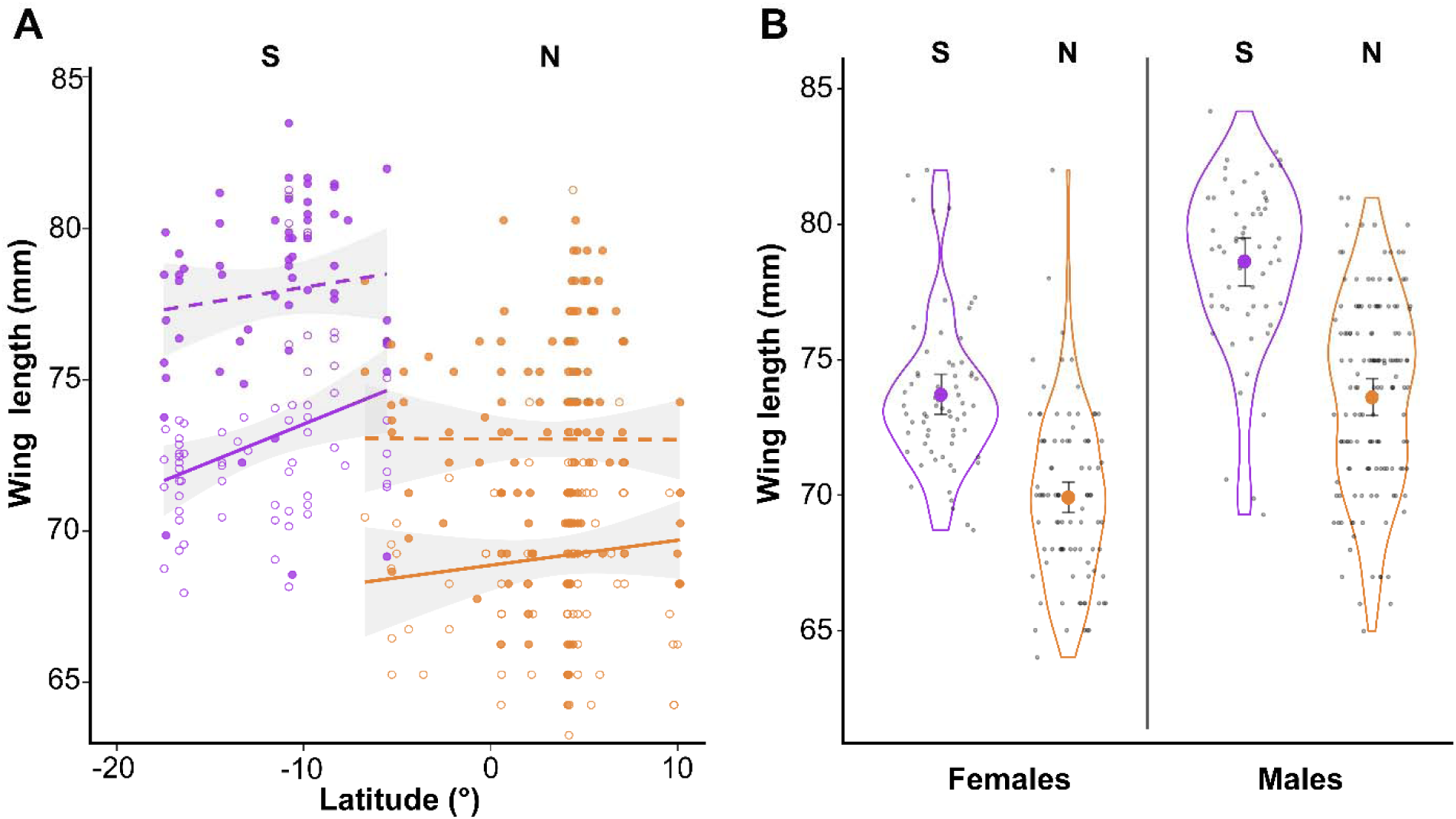
Geographic variation in wing length in *Diglossa cyanea* along its latitudinal range along the Andes, depicting the southern group (*D. c. melanopis*) in purple, and the northern group in orange. (A) Wing length does not show a range-wide latitudinal pattern of variation, although the southern group tended to have shorter wings towards its southern range limits. Females (open circles, solid fit line) and males (filled circles, dashed fit line) are indicated. (B) Both females and males of *D. c. melanopis*, south of the NPL (in purple) tended to be larger than individuals of their respective sex in the northern group (in orange). Despite the high variance in wing length and partial overlap between groups, linear models distinguish both the southern and northern groups based on wing-length data.

## DISCUSSION

By integrating genetic, vocal, and morphological data, we documented the existence of two divergent lineages within the Masked Flowerpiercer (*Diglossa cyanea*). Despite the extremely subtle plumage differences between these two lineages, they exhibit deep genetic structure on par with the amount of genetic divergence between other blue-plumaged flowerpiercer species in the *Diglossopis* clade. In addition, the two lineages have distinct songs in terms of structure and spectral traits, and dissimilar wing lengths in both males and females. Our results suggest two biological species in *D. cyanea* separated by the Northern Peruvian Low, as follows: *D. melanopis* from Peru and Bolivia, and *D. cyanea sensu stricto* from extreme northern Peru, Ecuador, Colombia, and Venezuela, including the subspecies *cyanea*, *dispar, obscura*, and *tovarensis*. Further evidence is needed to determine the degree of genetic, vocal, and phenotypic differentiation of *D. c. obscura* of the Sierra de Perijá, and *D. c. tovarensis* of the Coastal Cordillera of Venezuela.

The diversity within *Diglossa* has been likely shaped by time and geographical isolation, accompanied by genetic divergence, and possibly adaptive divergence in relation to environmental or social and behavioral factors (Moynihan 1979) that strengthen the effect of physical barriers (Vuilleumier 1984; Smith et al. 2014; Gutiérrez-Zuluaga et al. 2021). We have shown that in *D. cyanea,* the abrupt topographic and environmental turnover of the humid montane forest belt at the NPL maintains two diverging populations isolated since at least the Pliocene. Further, the lack of gene flow indicated by the sharp genetic break between the northern and southern populations of *D. cyanea* suggests the evolution of differences in traits, such as song, that could maintain separate lineages even when environmental changes permitted increased connectivity between populations, for example, by shifting forest belts during glacial periods (Hooghiemstra and Van der Hammen 2004; Ramírez-Barahona and Eguiarte 2013; Flantua and Hooghiemstra 2018).

Genetic divergence between the southern *D. cyanea* lineage (*D. c. melanopis*) was stronger with respect to northern *D. cyanea* (*sensu stricto*) than to *D. caerulescens,* and the ND2 gene tree showed a non-monophyletic *D. cyanea* as currently recognized. Although our genetic sampling encompassed most of the relevant geographic regions along the *D. cyanea* range, a few caveats in this phylogenetic inference should be noted. First, this gene tree may not reflect the true species history between these three taxa (Funk and Omland 2003; McKay and Zink 2010), so that a larger genetic dataset would be necessary to assess whether this phylogenetic hypothesis holds, especially given the extreme similarity in phenotype between the two *cyanea* groups. A plausible hypothesis to be tested is that the northern *D. cyanea* group and the southern *D. c. melanopis* are indeed sister species that originated from vicariance, or from dispersal over the NPL with subsequent differentiation, and this speciation event occurred shortly after the stem separation from the lineage leading to *D. caerulescens*. An approximation to address this first point would consist of vocal analyses that include *D. caerulescens* songs in the pairwise comparisons, which was beyond the scope of this study. A qualitative inspection of audio recordings of *D. caerulescens* songs indicates geographic variation across its range. Songs of *D. caerulescens* north of the NPL tend to be more similar to songs of the northern *D. cyanea* group, whereas south of the NPL *D. c. melanopis* and *D. caerulescens* seem much less similar. Differences in the bandwidth and duration of song sections are worthy of further research.

Another issue to be addressed is whether the absence of phenotypic or genetic samples of *Diglossa cyanea obscura*, of which few specimens exist (Zimmer and Phelps 1952), affects the analysis. Endemic to the Sierra de Perijá, this taxon is rare there (Hilty 2003; López-O et al. 2014), in contrast to the abundance of other subspecies of *Diglossa cyanea* within their ranges. This hints at ecological differences that might represent an additional cause of speciation. Finally, based on our limited genetic and vocal samples, we found that *D. c. tovarensis* is indeed a distinct population from the nominate *D. cyanea* of the Venezuelan and Colombian Andes; however, without additional data, we refrain from making additional taxonomic recommendations. It would be insightful to test experimentally via song playback how *tovarensis* reacts to *D. c. cyanea* songs.

In the oscine passerines, in addition to cultural evolution, plasticity in songs is common due to variation in learning abilities among individuals (Slater 1989; Price 2008); however, the genetically programmed, conservative template that predisposes learning of “own” species songs is informative of its phylogenetic history (Remsen 2005; Cadena and Cuervo 2010). In our study, the most significant differences in songs were found between the populations separated by the NPL. Most remarkably, the southern *D. c. melanopis* revealed two types of song, whereas the northern populations have only one. Additionally, the final whistling notes (S3), which are completely absent in the northern populations, may represent an innovation in song that is fixed in the southern *D. c*. *melanopis*, which in turn became its most distinctive vocal characteristic. Mirroring the genetic data, no vocal variation was observed within the large distribution of *D. c. melanopis*, or within that of the northern *D. cyanea* subspecies, except for the apparently distinct features of *D. c. tovarensis*. Freeman et al. (2022) found that *D. c. melanopis* discriminates between “own” songs and those of northern *D. c. cyanea*, and this reinforces the hypothesis that this level of song divergence would work as a reproductive barrier in case of secondary contact between these two lineages.

Our morphological analysis showed that wing length also shifts across the NPL. Although the only association with latitude was a slight decrease in WL towards the southern range limits, in Bolivia, of *D. c. melanopis*, the extent of sexual dimorphism in this trait is well conserved within *D. c. cyanea* and *D. c. melanopis* on both sides of the NPL (Fig. 6). Remarkably, WL differs on average, between the northern and southern populations, by more than 5 mm for males and 4 mm for females, representing an approximate 6% length difference between sexes based on the total WL.

In this study, we found evidence for an independent evolutionary history of the two *D. cyanea* lineages separated by the NPL that is expressed in unique vocal and phenotypic traits. Despite the conspicuous genetic, vocal, and size differences, minimal variation in plumage coloration and patterns have masked this diversity, as has been the case for multiple tropical birds (e.g. D’Horta et al. 2013; Smith et al. 2018; Berv et al. 2021). The evolutionary assembly of the Neotropical montane avifauna still has much to understand considering the complexity and diversity of evolutionary histories and the wide range of phenotypic divergences between populations (Weir 2009; Fjeldså et al. 2012; Cuervo 2013; Winger and Bates 2015; Cadena et al. 2020). In flowerpiercers, patterns of speciation and phenotypic evolution within the core *Diglossa* clade (Mauck and Burns 2009; Gutiérrez-Zuluaga et al. 2021), in which marked plumage diversity is evident but vocal variation is slight (pers. obs.), contrasts with our findings in the *Diglossopis* clade. Our results demonstrate that analyzing multiple characters reveal that the ecological and evolutionary patterns in *Diglossa* flowerpiercers are far more complex than previously recognized.

### Taxonomic implications

Based on our results, and after inspecting a large series of specimens, we recommend elevating the southern group to species rank: *Diglossa melanopis* (von Tschudi, 1844), with the English name Inca Flowerpiercer. No phenotypic variation has ever been recognized across its range, and we found no indication otherwise, rendering *D. melanopis* as a monotypic species. Therefore, *Diglossa cyanea* (Lafresnaye, 1840) may be restricted to all populations north and west of the North Peruvian Low. We recommended a change in its English name to Fire-eyed Flowerpiercer. We conclude that the subspecies *dispar* should not be recognized and be treated as a junior synonym of the nominate *D. cyanea* because specimens are undistinguishable. The number of available specimens of the much rarer Perijá population (*obscura*) is limited to assess its diagnosbility, and it may be better be subsumed with *cyanea*. In sum, *Diglossa cyanea* sensu stricto (Lafresnaye, 1840) is restricted to all populations north and west of the North Peruvian Low, with three subspecies: *cyanea*, *obscura*, and *tovarensis*.

## ACKNOWLEDGEMENTS

First, we thank the heroic efforts of field collectors, curators, genetic resources collections, museums, and sound recordists that make studies likes this possible. In particular, Louisiana State University Museum of Natural Science (D. Dittmann, S. Cardiff), Instituto de Ciencias Naturales, Universidad Nacional de Colombia (F. G. Stiles, N. Pérez), Instituto Alexander von Humboldt (D. López, S. Sierra, S. Pérez), Colección Ornitológica Phelps (M. Lentino), Academy of Natural Sciences at Drexel University (N. Rice, J. Weckstein), American Museum of Natural History (J. Cracraft, P. Sweet, T. J. Trombone), Field Museum (J. Bates, D. Willard), University of Kansas Natural History Museum (M. Robbins), Smithsonian National Museum of Natural History (G. Graves, J. Dean), Universidad Central de Venezuela (J. Pérez-Emán), Macaulay Library, Cornell Lab of Ornithology (G. Budney and M. Medler), xeno-canto (R. Planqué and W.-P. Vellinga). For support in the field, the collection, or the molecular laboratory we thank J. P. López, J. Pérez-Emán, J. Botero, J. Miranda, Y. López Padrón, J. Márquez, S. Sierra, J. E. Avendaño, K. Certuche, and G. Suárez. We thank C. Zábala and A. Cornejo for allowing us to use their photographs. This manuscript was improved thanks to comments by G. Knafler, T. S. Sillett and two anonymous reviewers. Special thanks to F. J. Urrea-Barreto who supported the study through its process and provided valuable comments for the manuscript. This study was partially funded by the Lewis and Clark Exploration Fund, Society of Systematic Biologists, Society of Integrative and Comparative Biology, F. M. Chapman Memorial Fund, American Ornithological Society, Wilson Ornithological Society, Idea Wild, and by National Science Foundation DDIG grants DEB-0910285

## SUPPLEMENTAL TABLES (headings)

**Supplementary Material Table S1.** Locality and collection information of all *Diglossa* tissue samples used in this study. AMNH = American Museum of Natural History; ANSP = Academy of Natural Sciences of Philadelphia; COP-IZET = Colección Ornitológica Phelps - Instituto de Zoología y Ecología Tropical; FMNH = Field Museum of Natural History; IAvH-CT = tissue collection of Instituto Alexander von Humboldt; ICN: Instituto de Ciencias Naturales, Universidad Nacional de Colombia; LSUMZ-B = Louisiana State University, Museum of Natural Science; MUA-AVP = Museo Universitario, Universidad de Antioquia; USNM-B = National Museum of Natural History, Smithsonian Institution.

**Supplementary Material Table S2.** Locality, sound library information and group assigned in analyses of all *Diglossa cyanea* song recording samples used in this study. ML = Macaulay Library, Cornell Lab of Ornithology; XC = xeno-canto.

**Supplementary Material Table S3.** Structure for all the Bayesian linear mixed models (BLMMs) used to assess whether vocal variables (duration, peak frequency, maximum frequency, minimum frequency, and bandwidth) of songs in the sections analyzed (F = Full song, S1 = Section 1, S2 = Section 2) predicted divergence in populations across the Northern Peruvian Low (NPL).

**Supplementary Material Table S4.** Locality, collection, age, sex, weight, wing length and group assigned in analyses of all *Diglossa cyanea* museum skin samples used in this study.

ANSP = Academy of Natural Sciences of Drexel University; COP = Colección Ornitológica Phelps; ICN = Instituto de Ciencias Naturales, Universidad Nacional de Colombia; MLS = Museo de la Salle, Bogotá; IAvH-A = Instituto Alexander von Humboldt; LSUMZ = Louisiana State University, Museum of Natural Science; MLZ: Moore Laboratory of Zoology, Occidental College.

**Supplementary Material Table S5.** Output from the Bayesian linear mixed models (BLMMs) that assesses whether vocal variables (duration, peak frequency, maximum frequency, minimum frequency, and bandwidth) of songs in the sections analyzed (F = Full song, S1 = Section 1, S2 = Section 2) predicted divergence in populations across the Northern Peruvian Low (NPL). Regression coefficients (β) and variance components (σ2) are reported with the 95% Confidence Interval (CI). Values are presented in their original scale: seconds (s) for Duration, and Hertz (Hz) for frequency variables. We present highlighted statistically significant difference in all five variables. Sample size as follows: n_[recordings]_ = 81, n_[song bouts]_ = 325, n_[northern group recordings]_ = 53, n_[southern group recordings]_ = 28.

**Supplementary Material Table S6.** Output from the Bayesian linear mixed models (BLMMs) that assesses whether vocal variables (duration, peak frequency, maximum frequency, minimum frequency, and bandwidth) of songs in the sections analyzed (F = Full song, S1 = Section 1, S2 = Section 2) predicted divergence in populations across the Northern Peruvian Low (NPL). Regression coefficients (β) and variance components (σ2) are reported with the 95% Confidence Interval (CI). Values are presented in a latent (standardized) scale. We present highlighted statistically significant difference in all five variables. Sample size as follows: n_[recordings]_ = 81, n_[song bouts]_ = 325, n_[northern group recordings]_ = 53, n_[southern group recordings]_ = 28.

**Supplementary Material Table S7.** Output from the Bayesian linear mixed models (BLMMs) that assesses whether vocal variables (duration, peak frequency, maximum frequency, minimum frequency, and bandwidth) of songs in the sections analyzed (F = Full song, S1 = Section 1, S2 = Section 2) predicted differences between *D. cyanea* subspecies. Regression coefficients (β) and variance components (σ2) are reported with the 95% Confidence Interval (CI). Values are presented in their original scale. We present highlighted statistically significant difference in all five variables. Sample size as follows: n_[recordings]_ = 81, n_[song bouts]_ = 325, n_[*melanopis* recordings]_ = 28, n_[*dispar* group recordings]_ = 6, n_[*cyanea* recordings]_ = 45, n_[*tovarensis* recordings]_ = 2.

**Supplementary Material Table S8.** Outputs from the Bayesian linear mixed models (BLMMs) that assess whether vocal variables (duration, peak frequency, maximum frequency, minimum frequency, and bandwidth) of the full song (F) predicted differences between alternative songs in the southern populations and the main songs at both sides of the NPL. Regression coefficients (β) and variance components (σ^2^) are reported with the 95% Confidence Interval (CI). Values are presented in their original scale. We present highlighted statistically significant difference in all five variables. Sample size as follows: n_[recordings]_ = 88, n_[song bouts]_ = 382, n_[alternative song recordings]_= 16, n_[northern son recordings]_ = 53, n_[main southern song recordings]_ = 28. Note: 18 of the recordings analyzed contained both main and alternative southern songs.

**Supplementary Material Table S9.** Outputs from the linear models (LMs) that assess differences in wing length across a latitude gradient separated by sex and analyzed separately from northern and southern populations to the NPL. Sample size as follows: n_[northern females]_ = 95, n_[Southern females]_ = 66, n_[northern males]_ = 144, n_[southern males]_ = 59.

**Supplementary Material Table S10.** Outputs from the linear models (LMs) that assess differences in wing length between males from northern and southern populations and between females from northern and southern groups. Sample size as follows: n_[northern females]_ = 95, n_[southern females]_ = 66, n_[northern males]_ = 144, n_[southern males]_ = 59.

## SUPPLEMENTAL FIGURES WITH CAPTIONS

**Supplementary Material Figure S1.**
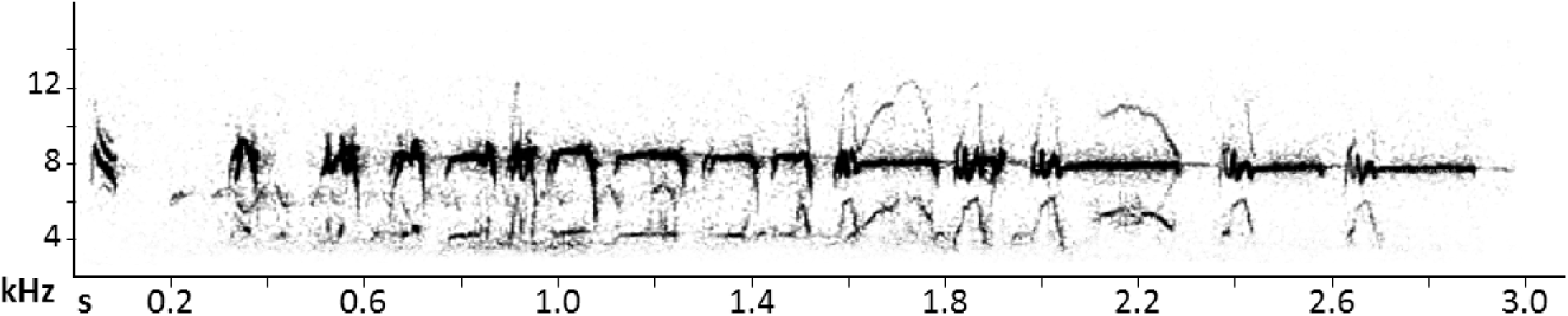
Spectrogram of a representative sample of the alternative song in the southern populations of Masked Flowerpiercer, *D. c. melanopis* (from Pasco, Peru, ML 35825). Note that the general structure of the typical or main song is not followed here. While this song begins with few short “*tzi*” notes, there are not well differentiated S2 and S3 sections; the latter half of the song are strident whistling notes (each of them with a little “*tzi*” within them) that become longer as the song ends.

**Supplementary Material Figure S2.**
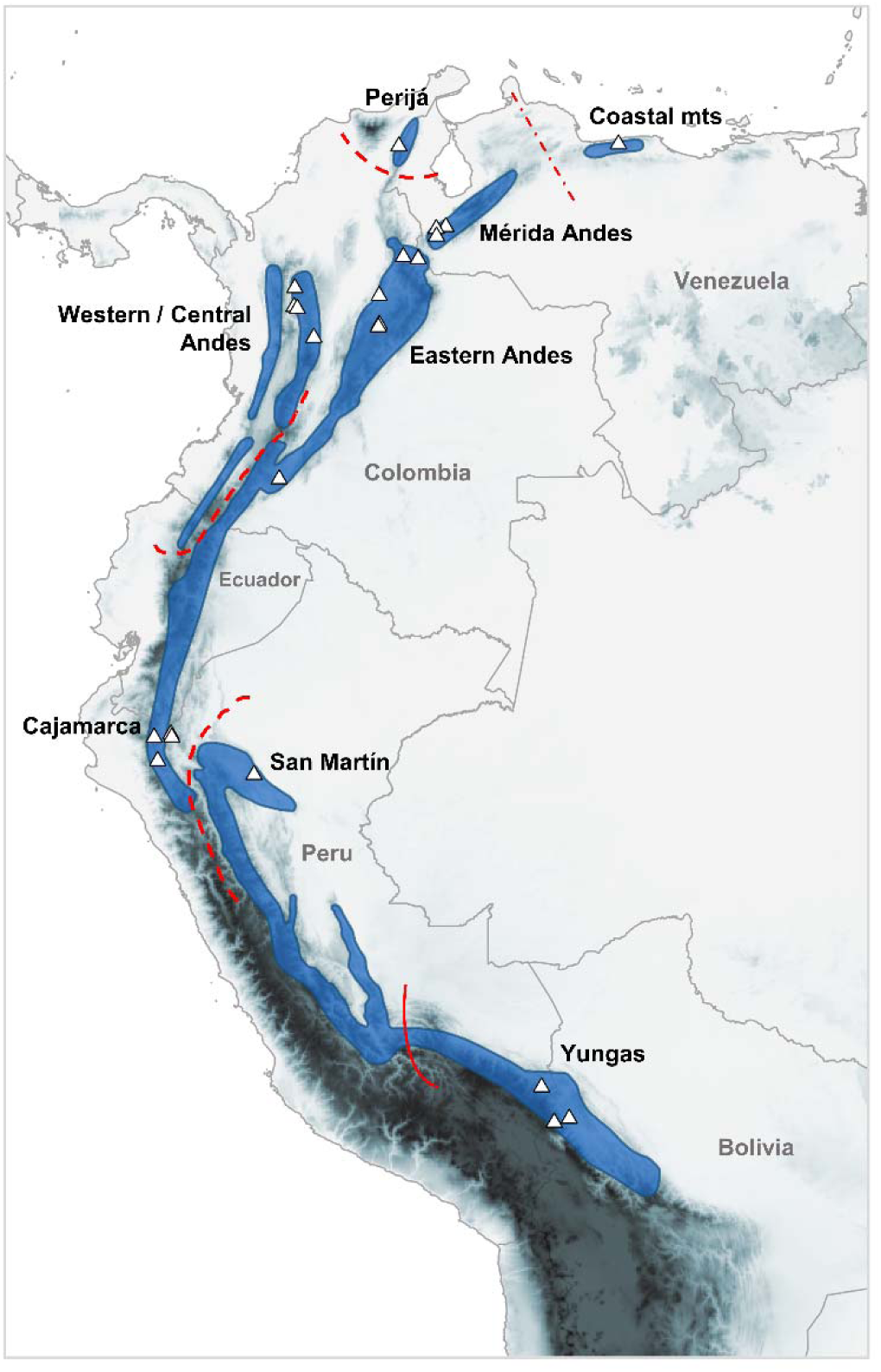
Geographic distribution of the Bluish Flowerpiercer, *Diglossa caerulescens*. Red lines indicate approximate range boundaries between the divergent populations according to our phylogeographic analysis. Subspecies range boundaries usually coincide with low elevation gaps. White triangles represent sample localities for vouchered specimens of *D. caerulescens* used in the phylogeographic analysis.

**Supplementary Material Figure S3.**
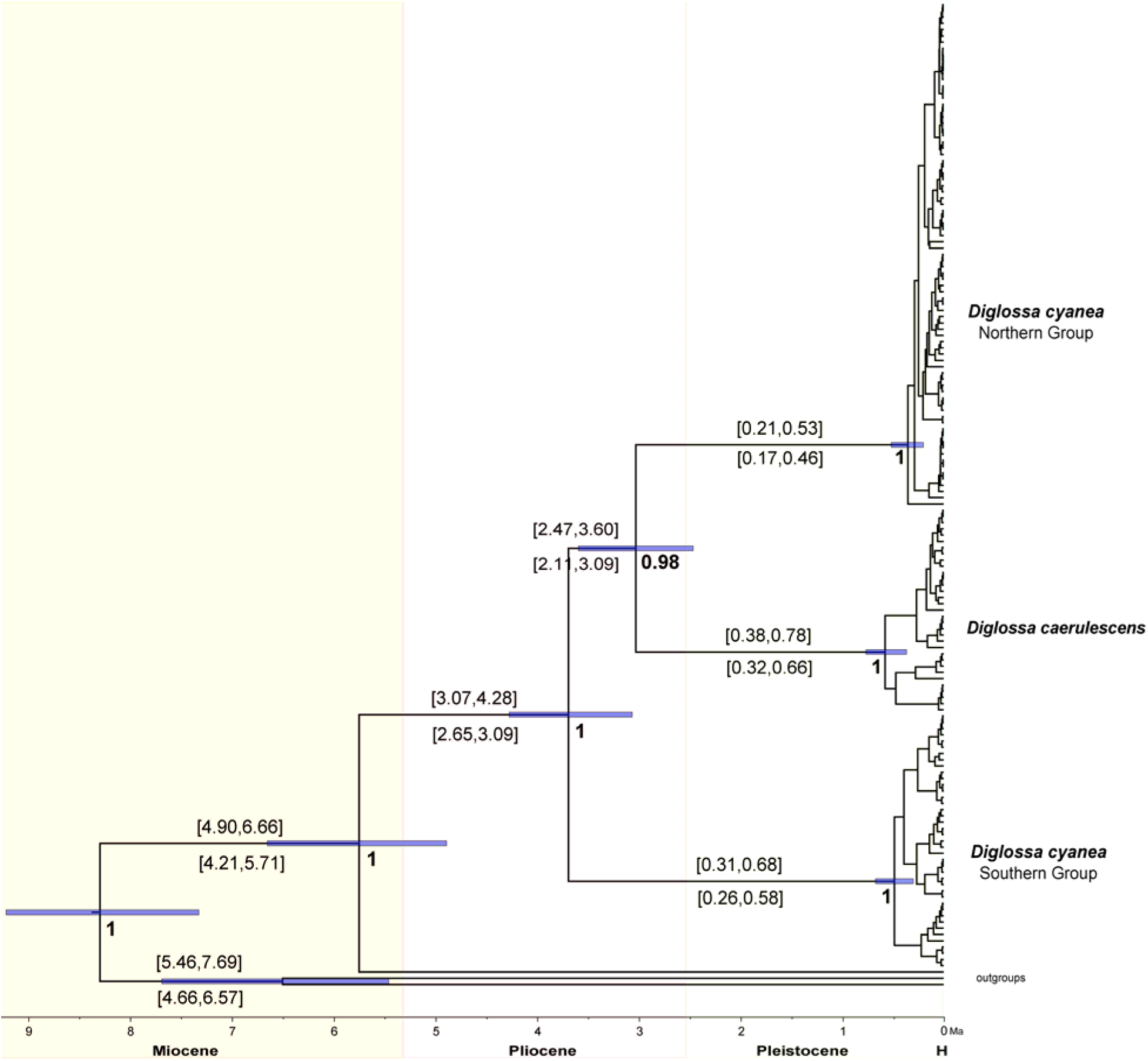
BEAST maximum clade credibility tree based on ND2 data for all individuals sampled, showing two diverging lineages in *D. cyanea*: the northern group (including subspecies *cyanea*, *dispar*, and *tovarensis*), and the southern group (subspecies *melanopis*). Similar to the MrBayes tree topology (Fig. 3), this analysis indicates that *D. cyanea* is not monophyletic because *D. caerulescens* is the sister lineage to the northern *D. cyanea* group. The two speciation events leading to these three lineages date back to the Late Pliocene, and largely overlap in time between them. Node bars and range values on top of branches (in Ma), report the 95% highest posterior density intervals on divergence times using the 2.5% rate. Range values below branches report the estimated divergence times using the faster rate of 2.9% (see text). Values next to nodes are the posterior probabilities, which were identical for both analyses (with the two rates).

**Supplementary Material Figure S4.**
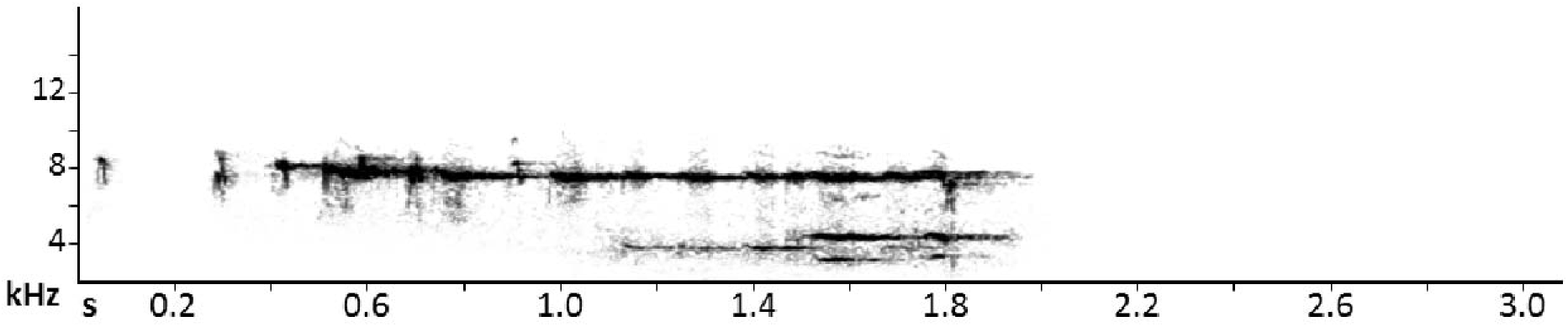
Spectrogram sample of the song of *D. c. tovarensis* populations (from Aragua, northern Venezuela, ML 222433). Note that it is possible to identify S1 and S2 as in the other typical songs, but note shape is less conspicuous, bandwidth is relatively narrower, with lower maximum frequency and shorter S1 duration than in all other northern subspecies (*cyanea, dispar*).

## LITERATURE CITED

Bandelt, H. J., P. Forster, and A. Rohl (1999). Median-joining networks for inferring intraspecific phylogenies. Molecular Biology and Evolution 16:37–48.

Benham, P. M., A. M. Cuervo, J. A. McGuire, and C. C. Witt (2014). Biogeography of the Andean metaltail hummingbirds: Contrasting evolutionary histories of tree line and habitat-generalist clades. Journal of Biogeography 42:763–777.

Berv, J. S., L. Campagna, T. J. Feo, I. Castro-Astor, C. C. Ribas, R. O. Prum, and I. J. Lovette (2021). Genomic phylogeography of the White-crowned Manakin *Pseudopipra pipra* (Aves: Pipridae) illuminates a continental-scale radiation out of the Andes. Molecular Phylogenetics and Evolution 164:107205.

Bock, W. J. (1985). Is *Diglossa* (Thraupinae) monophyletic? Ornithological Monographs 36:319–332.

Bolker, B. M., M. E. Brooks, C. J. Clark, S. W. Geange, J. R. Poulsen, M. H. H. Stevens, and J. S. S. White (2009). Generalized linear mixed models: a practical guide for ecology and evolution. Trends in Ecology and Evolution 24:127–135.

Bouckaert, R., T. G. Vaughan, J. Barido-Sottani, S. Duchêne, M. Fourment, A. Gavryushkina, J. Heled, G. Jones, D. Kühnert, N. De Maio, M. Matschiner, et al. (2019). BEAST 2.5: An advanced software platform for Bayesian evolutionary analysis. PLoS Computational Biology 15:e1006650.

Cadena, C. D., and A. M. Cuervo (2010). Molecules, ecology, morphology, and songs in concert: how many species is *Arremon torquatus* (Aves: Emberizidae)? Biological Journal of the Linnean Society 99:152–176.

Cadena, C. D., J. L. Pérez-Emán, A. M. Cuervo, L. N. Céspedes, K. L. Epperly, and J. T. Klicka (2019). Extreme genetic structure and dynamic range evolution in a montane passerine bird: implications for tropical diversification. Biological Journal of the Linnean Society 126:487–506.

Cadena, C. D., A. M. Cuervo, L. N. Céspedes, G. A. Bravo, N. Krabbe, T. S. Schulenberg, G. E. Derryberry, L. F. Silveira, E. P. Derryberry, R. T. Brumfield, and J. Fjeldså (2020). Systematics, biogeography, and diversification of *Scytalopus* tapaculos (Rhinocryptidae), an enigmatic radiation of Neotropical montane birds. Auk 137:1–30.

Chaves, J. A., J. T. Weir, and T. B. Smith (2011). Diversification in *Adelomyia* hummingbirds follows Andean uplift. Molecular Ecology 20:4564–4576.

Coyne, J. A., and H. A. Orr. 2004. Speciation. 1 edition. Sinauer Associates, Sunderland, MA, USA.

Cracraft, J. (1985). Historical biogeography and patterns of differentiation within the South American avifauna: Areas of endemism. Ornithological Monographs 36:49–84.

Cuervo, A. M. (2013). Evolutionary Assembly of the Neotropical Montane Avifauna. Ph.D. Dissertation, Louisiana State University, Baton Rouge, LA, USA.

Cuervo, A. M., F. G. Stiles, M. Lentino, R. T. Brumfield, and E. P. Derryberry (2014). Geographic variation and phylogenetic relationships of *Myiopagis olallai* (Aves: Passeriformes; Tyrannidae), with the description of two new taxa from the Northern Andes. Zootaxa 3873:1–24.

D’Horta, F. M., A. M. Cuervo, C. C. Ribas, R. T. Brumfield, and C. Y. Miyaki (2013). Phylogeny and comparative phylogeography of *Sclerurus* (Aves: Furnariidae) reveal constant and cryptic diversification in an old radiation of rain forest understorey specialists. Journal of Biogeography 40:37–49.

Dickinson, E. C., and L. Christidis. 2014. The Howard & Moore Complete Checklist of the Birds of the World. 4th. edition, Vol. 2: Passerines. 4th. edition, Vol. 2 edition. Aves Press, Eastbourne, United Kingdon.

Duellman, W. E. (1979). The herpetofauna of the Andes: Patterns of distribution, origin, differentiation, and present communities. In The South American Herpetofauna: Its Origin, Evolution, and Dispersal, vol. 7 (W. E. Duellman, Editor). Monograph of the Museum of Natural History, University of Kansas, Lawrence, KS, USA:371–459.

Fjeldså, J., and N. Krabbe. 1990. Birds of the High Andes. Zoological Museum, University of Copenhagen and Apollo Books, Svendborg, Denmark.

Fjeldså, J., R. C. K. Bowie, and C. Rahbek (2012). The Role of Mountain Ranges in the Diversification of Birds. Annual Review of Ecology, Evolution, and Systematics 43:249–265.

Flantua, S. G., and H. Hooghiemstra (2018). Historical connectivity and mountain biodiversity. In Mountains, climate and biodiversity (C. Hoorn, A. Perrigo, and A. Antonelli, Editors). John Wiley & Sons Ltd:171–185.

Freeman, B. G., J. Rolland, G. A. Montgomery, and D. Schluter (2022). Faster evolution of a premating reproductive barrier is not associated with faster speciation rates in New World passerine birds. Proceedings of the Royal Society B 289:1–8.

Freile, J. F., and R. Restall. 2018. Birds of Ecuador. Bloomsbury Publishing, New York, NY, USA.

Funk, D. J., and K. E. Omland (2003). Species-level paraphyly and polyphyly: Frequency, causes, and consequences, with insights from animal mitochondrial DNA. Annual Review of Ecology, Evolution, and Systematics 34:397–423.

Graves, G. R. (1982). Speciation in the Carbonated Flower-Piercer (*Diglossa carbonaria*) complex of the Andes. Condor 84:1–14.

Graves, G. R. (1985). Elevational correlates of speciation and intraspecific geographic variation in plumage in Andean forest birds. Auk 102:556–579.

Graves, G. R. (1988). Linearity of geographic range and its possible effect on the population structure of Andean birds. Auk 105:47–52.

Graves, G. R. (1991). Bergmann’s rule near the equator: Latitudinal clines in body size of an Andean passerine bird. Proceedings of the National Academy of Sciences of the United States of America 88:2322–2325.

Gutiérrez-Pinto, N., A. M. Cuervo, J. Miranda, J. L. Pérez-Emán, R. T. Brumfield, and C. D. Cadena (2012). Non-monophyly and deep genetic differentiation across low-elevation barriers in a Neotropical montane bird (*Basileuterus tristriatus*; Aves: Parulidae). Molecular Phylogenetics and Evolution 64:156–165.

Gutiérrez-Pinto, N., K. G. McCracken, P. L. Tubaro, C. Kopuchian, A. Astie, and C. D. Cadena (2019). Molecular and morphological differentiation among Torrent Duck (*Merganetta armata*) populations in the Andes. Zoologica Scripta 48:589–604.

Gutiérrez-Zuluaga, A. M., C. González-Quevedo, J. A. Oswald, R. S. Terrill, J. L. Pérez-Emán, and J. L. Parra (2021). Genetic data and niche differences suggest that disjunct populations of *Diglossa brunneiventris* are not sister lineages. Ornithology 138:1–14.

Hadfield, J. D. (2010). MCMC methods for multi-response generalized linear mixed models: The MCMCglmm R package. Journal of Statistical Software 33:1–22.

Heled, J., and A. J. Drummond (2010). Bayesian inference of species trees from multilocus data. Molecular Biology and Evolution 27:570-580.

Hellmayr, C. E. (1935). Catalogue of Birds of the Americas and the Adjacent Islands, Vol. 13, Part VIII. Field Museum of Natural History Zoological Series 13.

Herzog, S. K., R. S. Terrill, A. E. Jahn, J. J. V. Remsen, O. M. Z, V. H. García-Solíz, R. MacLeod, A. MacCormick, and J. Q. Vidoz. 2017. Birds of Bolivia.

Hiller, A. E., R. T. Brumfield, and B. C. Faircloth (2021). A reference genome for the nectar-robbing Black-throated Flowerpiercer (*Diglossa brunneiventris*). G3 Genes|Genomes|Genetics 11.

Hilty, S. L. 2003. Birds of Venezuela. Second edition. Princeton University Press, New Jersey, NJ.

Hooghiemstra, H., and T. Van der Hammen (2004). Quaternary ice-age dynamics in the Colombian Andes: developing an understanding of our legacy. Philosophical Transactions of the Royal Society of London. Series B: Biological Sciences 359:173–181.

Isler, M. L., and P. R. Isler. 1999. The Tanagers: Natural history, Distribution, and Identification. Second edition. Smithsonian Institution Press, Washington, D. C.

Isler, M. L., R. T. Chesser, M. B. Robbins, A. M. Cuervo, C. D. Cadena, and P. A. Hosner (2020). Taxonomic evaluation of the *Grallaria rufula* (Rufous Antpitta) complex (Aves: Passeriformes: Grallariidae) distinguishes sixteen species. Zootaxa 4817:1–74.

Krabbe, N., and T. S. Schulenberg (1997). Species limits and natural history of *Scytalopus* tapaculos (Rhinocryptidae), with descriptions of the Ecuadorian taxa, including three new species. Ornithological Monographs 48:47–88.

Kumar, S., G. Stecher, M. Li, C. Knyaz, and K. Tamura (2018). MEGA X: Molecular evolutionary genetics analysis across computing platforms. Molecular Biology and Evolution 35:1547–1549.

Lanfear, R., P. B. Frandsen, A. M. Wright, T. Senfeld, and B. Calcott (2017). Partitionfinder 2: New methods for selecting partitioned models of evolution for molecular and morphological phylogenetic analyses. Molecular Biology and Evolution 34:772–773.

Leigh, J. W., and D. Bryant (2015). PopART: Full-feature software for haplotype network construction. Methods in Ecology and Evolution 6:1110–1116.

López-O, J. P., J. E. Avendaño, N. Gutiérrez-Pinto, and A. M. Cuervo (2014). The birds of the serranía de Perijá: The northernmost avifauna of the Andes. Ornitologia Colombiana 14:62–93.

Mauck, W. M., and K. J. Burns (2009). Phylogeny, biogeography, and recurrent evolution of divergent bill types in the nectar-stealing flowerpiercers (Thraupini: *Diglossa* and *Diglossopis*). Biological Journal of the Linnean Society 98:14–28.

Mayr, E. 1942. Systematics and the Origin of Species. 1 edition, New York, NY, USA.

Mayr, E. 1963. Animal Species and Evolution. 1 edition, Cambridge, MA, USA.

McKay, B. D., and R. M. Zink (2010). The causes of mitochondrial DNA gene tree paraphyly in birds. Molecular Phylogenetics and Evolution 54:647–650.

Meyer de Schauensee, R. (1951). The birds of the Republic of Colombia (Cuarta entrega: Alaudidae-Fringillidae). Caldasia 5:873–1112.

Miller, M. A., W. Pfeiffer, and T. Schwartz. 2010. Creating the CIPRES Science Gateway for inference of large phylogenetic trees. Pages 1-8 in 2010 Gateway Computing Environments Workshop, GCE 2010, New Orleans, LA, U.S.A.

Moynihan, M. (1968). Social mimicry; character convergence versus character displacement. Evolution 22:315–331.

Moynihan, M. (1979). Geographic variation in social behavior and in adaptations to competition among Andean birds. Publications of the Nuttall Ornithological Club 18:1.

Nakagawa, S., and I. C. Cuthill (2007). Effect size, confidence interval and statistical significance: A practical guide for biologists. Biological Reviews 82:591–605.

Nosil, P. (2008). Ernst Mayr and the integration of geographic and ecological factors in speciation. Biological Journal of the Linnean Society 95:26–46.

Núñez-Zapata, J., L. E. Pollack-Velásquez, E. Huamán, J. Tiravanti, and E. García (2016). A compilation of the birds of La Libertad Region, Peru. Revista Mexicana de Biodiversidad 87:200–215.

Palacios, C., S. García-R, J. L. Parra, A. M. Cuervo, F. G. Stiles, J. E. McCormack, and C. D. Cadena (2019). Shallow genetic divergence and distinct phenotypic differences between two Andean hummingbirds: Speciation with gene flow? The Auk 136:1–21.

Parker, T. A., III, D. F. Stotz, and J. W. Fitzpatrick (1996). Ecological and distributional databases for Neotropical birds. In Neotropical birds: Ecology and conservation (D. F. Stotz, J. W. Fitzpatrick, T. A. Parker, III, and D. Moskovits, Editors). Chicago University Press, Chicago.

Parker, T. A., III, T. S. Schulenberg, G. R. Graves, and M. J. Braun (1985). The avifauna of the Huancabamba region, northern Peru. Ornithological Monographs 36:169–197.

Patten, M. A. (2010). Null expectations in subspecies diagnosis. Ornithological Monographs 67:35–41.

Paynter, R. A. 1981. Ornithological gazetteer of Colombia. 2nd edition. Museum of Comparative Zoology, Harvard University, Cambridge, MA, USA.

Paynter, R. A. 1992. Ornithological gazetteer of Bolivia. 2nd edition, Cambridge, MA, USA.

Price, T. D. 2008. Speciation in Birds. 1 edition. Roberts & Company, Greenwood Village, CO, USA.

Prieto-Torres, D. A., A. M. Cuervo, and E. Bonaccorso (2018). On geographic barriers and Pleistocene glaciations: Tracing the diversification of the Russet-crowned Warbler (*Myiothlypis coronata*) along the Andes. PLoS ONE 13:e0191598.

R Core Team (2021). R: A Language and Environment for Statistical Computing.

Rambaut, A., A. J. Drummond, D. Xie, G. Baele, and M. A. Suchard (2018). Posterior summarization in bayesian phylogenetics using Tracer 1.7. Systematic Biology 67:901–904.

Ramírez-Barahona, S., and L. E. Eguiarte (2013). The role of glacial cycles in promoting genetic diversity in the Neotropics: the case of cloud forests during the Last Glacial Maximum. Ecology and Evolution 3:725–738.

Remsen, J. V., Jr. (1984). High incidence of “leapfrog” pattern of geographic variation in Andean Birds: Implications for the speciation process. Science 224:171–173.

Remsen, J. V., Jr. (2005). Pattern, process, and rigor meet classification. Auk 122:403–413.

Remsen, J. V., Jr. (2010). Subspecies as a meaningful taxonomic rank in avian classification. Ornithological Monographs 67:62–78.

Ronquist, F., M. Teslenko, P. Van Der Mark, D. L. Ayres, A. Darling, S. H. Ohna, B. Larget, L. Liu, M. A. Suchard, and J. P. Huelsenbeck (2012). MrBayes 3.2: Efficient Bayesian Phylogenetic Inference and Model Choice Across a Large Model Space. Systematic Biology 61:539–542.

Schielzeth, H. (2010). Simple means to improve the interpretability of regression coefficients. Methods in Ecology and Evolution 1:103–113.

Schulenberg, T. S., D. F. Stotz, D. F. Lane, J. P. O’Neill, and T. A. Parker, III. 2010. Birds of Peru. Revised and Updated Edition edition. Princeton University Press, Princeton, NJ, USA.

Slater, P. J. B. (1989). Bird song learning: Causes and consequences. Ethology Ecology and Evolution 1:19–46.

Smith, B. T., and J. Klicka (2010). The profound influence of the Late Pliocene Panamanian uplift on the exchange, diversification, and distribution of New World birds. Ecography.

Smith, B. T., R. W. Bryson, W. M. Mauck, J. Chaves, M. B. Robbins, A. Aleixo, and J. Klicka (2018). Species delimitation and biogeography of the gnatcatchers and gnatwrens (Aves: Polioptilidae). Molecular Phylogenetics and Evolution 126:45–57.

Smith, B. T., J. E. McCormack, A. M. Cuervo, M. J. Hickerson, A. Aleixo, C. D. Cadena, J. Pérez-Emán, C. W. Burney, X. Xie, M. G. Harvey, B. C. Faircloth, et al. (2014). The drivers of tropical speciation. Nature 515:406–409.

Stamatakis, A. (2014). RAxML version 8: A tool for phylogenetic analysis and post-analysis of large phylogenies. Bioinformatics 30:1312–1313.

Stecher, G., K. Tamura, and S. Kumar (2020). Molecular evolutionary genetics analysis (MEGA) for macOS. Molecular Biology and Evolution 37:1237–1239.

Stephens, L., and M. A. Traylor. 1983. Ornithological Gazetteer of Peru, Cambridge, MA, USA.

Valderrama, E., J. L. Pérez-Emán, R. T. Brumfield, A. M. Cuervo, and C. D. Cadena (2014). The influence of the complex topography and dynamic history of the montane Neotropics on the evolutionary differentiation of a cloud forest bird (*Premnoplex brunnescens*, Furnariidae). Journal of Biogeography 41:1533–1546.

Vuilleumier, F. (1968). Population structure of the *Asthenes flammulata*, superspecies (Aves: Furnariidae). Breviora 297:1–21.

Vuilleumier, F. (1969). Systematics and evolution in *Diglossa* (Aves, Coerebidae). American Museum Novitates 2381:1–44.

Vuilleumier, F. (1984). Zoogeography of Andean birds: Two major barriers, and speciation and taxonomy of the *Diglossa carbonaria* superspecies. National Geographic Society Research Reports 16:713–731.

Weir, J. T. (2009). Implications of genetic differentiation in Neotropical montane forest birds. Annals of the Missouri Botanical Garden 96:410–433.

Winger, B. M., and J. M. Bates (2015). The tempo of trait divergence in geographic isolation: Avian speciation across the Marañón Valley of Peru. Evolution 69:772–787.

Zimmer, J. T. (1942). Notes on the genera *Diglossa* and *Cyanerpes*, with addenda to *Ochthoeca*. American Museum Novitates 1203:2–15.

Zimmer, J. T., and W. H. Phelps (1952). A new race of the honey-creeper, *Diglossa cyanea*, from Venezuela. American Museum Novitates 1603:1–2.

Zink, R. M., and J. V. Remsen, Jr. (1986). Evolutionary processes and patterns of geographic variation in birds. In Current Ornithology, Vol. 4, vol. 4 (R. F. Johnston, Editor). Plenum Press, New York, NY, USA:1–69.

